# Microcompartments in archaeal ancestors of eukaryotes: a bioenergetic engine that could have fuelled eukaryogenesis

**DOI:** 10.1101/2025.11.08.687404

**Authors:** Huan Du, Aoxiang Xu, Xiaoyuan Feng, Wen-cong Huang, Huayu Li, Li Liu, Yuling Li, Siyu Zhang, Ning Song, Kathryn E. Appler, Brett J. Baker, Eugene V. Koonin, Meng Li, Yang Liu

## Abstract

Eukaryotic intracellular compartmentalization is a key innovation in the evolution of complex cellular life. While microcompartments enable metabolic specialization in many bacteria, to our knowledge, no analogous systems have been identified in Archaea. Here, we report the discovery of archaeal microcompartments (AMCs) in Hodarchaeales, an order within the phylum *Promethearchaeati* (Asgard archaea) that includes the closest known archaeal relatives of eukaryotes. Phylogenetic and structural analyses indicate that these catabolic AMCs, which are specialized for sugar-phosphate metabolism, were acquired by horizontal gene transfer from deep-rooted bacteria of the phylum Myxococcota. The shell pentamers of AMCs are fused to lysine/arginine-rich intrinsically disordered regions that capture cytosolic DNA, facilitating nutrient scavenging. Reaction-diffusion modelling predicts that enzyme colocalization and substrate channelling within AMCs can increase the NADH flux approximately 100-fold. Thus, the AMCs substantially boost energy production in the cell and might have primed the archaeal host for eukaryogenesis.

## Introduction

The origin of eukaryotes is one of the most momentous evolutionary innovations in Earth’s biological history. Reconstruction of the evolutionary trajectory from the closest prokaryotic relatives of eukaryotes (First Eukaryotic Common Ancestor, FECA) to the Last Eukaryotic Common Ancestor (LECA) remains a fundamental challenge^1^. The discovery of Asgard archaea (now formally known as *Promethearchaeati*), which encode a broad diversity of eukaryotic signature proteins (ESPs) and are phylogenetically identified as the closest relatives of eukaryotes^2–4^, revolutionized the study of eukaryogenesis^5–8^. Phylogenomic analyses leave no reasonable doubt regarding the origin of eukaryotes from within the Asgard diversity, and the latest findings suggest that the FECA branched out either at the base of the Heimdallarchaeia class^9^ or within it, as a sister group to the order Hodarchaeales^4^. The FECA likely evolved increased intracellular complexity, including membrane remodelling and cytoskeleton formation, features supported by the ESPs and present in some Asgard archaea^2–4^. These developments could have provided benefits, in particular, facilitating acquisition of endosymbionts, while imposing energetic costs, although this trade-off remains disputed^10,11^.

To understand eukaryogenesis, it is essential to clarify how FECA overcame its bioenergetic challenges to sustain genomic complexity before acquiring mitochondria. The use of oxygen is a key strategy by which microorganisms enhance their energy-requiring activities^12^. Genomic and structural analyses suggest that FECA possessed adaptations for oxygen metabolism, including genes for electron transport chain complex IV, heme biosynthesis, and reactive oxygen species (ROS) detoxification^13^. However, a physiological study on the first isolated Hodarchaeales strain suggested that the products of these genes were primarily involved in oxygen detoxification rather than aerobic respiration for energy production, as no growth was observed under (micro)aerobic conditions^14^. A parallel study showed that some Asgard archaea, such as Hodarchaeales and Kariarchaeaceae, exhibit oxygen adaptation mediated by terminal oxidase and globin^15^. Taken together, these findings suggest that the FECA tolerated oxygen but most likely did not use aerobic respiration for enhanced energy generation and might have possessed alternative mechanisms for increased energy production efficiency.

Compartmentalization enabled by membrane-bound organelles is one of the fundamental evolutionary innovations distinguishing eukaryotic from prokaryotic cells. Although lacking *bona fide* membrane-bound organelles, many bacteria possess organelle-like microcompartments, which serve as devices for enhanced metabolic efficiency and energy generation^16^. These bacterial microcompartments (BMCs) consist of a semi-permeable proteinaceous shell that encapsulates a suite of specialized enzymes, forming a dedicated microenvironment that enhances the efficiency of specific metabolic pathways by concentrating the respective enzymes and substrates^17–20^. The protein shell of a BMC is a polyhedron, typically, an icosahedron, assembled from three main types of proteins: hexamers (BMC-H/H^p^) and pseudohexamers (BMC-T^s^/T^sp^/T^dp^, form the facets, whereas pentamers (BMC-P), occupy the vertices^16^. The BMC-T proteins are dimers of the hexamer-forming BMC-H. The icosahedral structure of the BMC resembles the capsid structures of viruses, such as tailless bacteriophages in the family *Tectiviridae*. However, protein structure comparisons detected no relationship between the structural proteins of the BMC and viral capsid proteins. Instead, BMC-H and BMC-P apparently evolved from cellular ancestors, namely, PII signaling protein and OB-fold domain-containing protein, respectively^21^. Recent comparative genomic analyses have documented the presence of BMCs in 45 bacterial phyla, revealing both anabolic types involved in carbon fixation^22^ and diverse catabolic variants specialized for the metabolism of aromatics, ethanol, or sugar-phosphate^23^. In contrast, to our knowledge, no evidence of the existence of microcompartments in archaea has been reported so far.

Here, we report the discovery of microcompartments (archaeal microcompartments, AMCs) in Hodarchaeales. We identified a pronounced enrichment of gene clusters encoding homologs of sugar phosphate utilization (SPU) microcompartment proteins across diverse Hodarchaeales lineages and demonstrated the self-assembly of the AMC shell. Phylogenetic analyses indicate that the Hodarchaeales AMC operons were horizontally acquired from a deep-branching Myxococcota lineage, with ancestral reconstruction tracing both AMC genes and oxygen-adaptation proteins to the last Hodarchaeales common ancestor (LHoCA). Mathematical modelling predicted that the AMC shell trapping could increase the NADH conversion rate more than 100-fold, potentially enabling high energy production. These findings are compatible with the possibility that AMCs functioned as a critical bioenergetic module in the FECA, illustrating how horizonal gene transfer (HGT) could have set the scene for the mitochondrial endosymbiosis and eukaryogenesis.

## Results

### Distribution of proteinaceous microcompartment gene clusters in Archaea domain

We analyzed more than 15,000 dereplicated representative archaeal genomes (see Methods, Supplementary Table 1). Comprehensive genomic screening identified putative microcompartment gene clusters across the Archaea domain. Homologs of the core shell proteins, BMC-H (BMC domain, PF00936, hexamer or trimer) and BMC-P (ethanolamine utilisation protein, EutN/carboxysome, PF03319, pentamer) were found to be broadly distributed in multiple archaeal lineages, including Asgard, DPANN (mainly SpSt-1190), and TACK superphyla (mainly JASLWR01 order from Thermo-plasmatota), displaying a patchy distribution in each of these groups (Fig. 1, Supplementary Fig. 1). Microcompartment gene clusters were detected in 66 of the 111 analyzed Hodarchaeales genomes^13^, showing a much higher prevalence and genetic diversity (total branch length of 7.38, a proxy for phylogenetic diversity; see Methods) compared to other archaeal lineages (e.g., 0.08 for SpSt-1190 and 0.93 for JASLWR01). In particular, the AMC operon is present in the isolated Hodarchaeales strain SC1, one of the only two cultured representatives of this order^14^.

**Figure 1:**
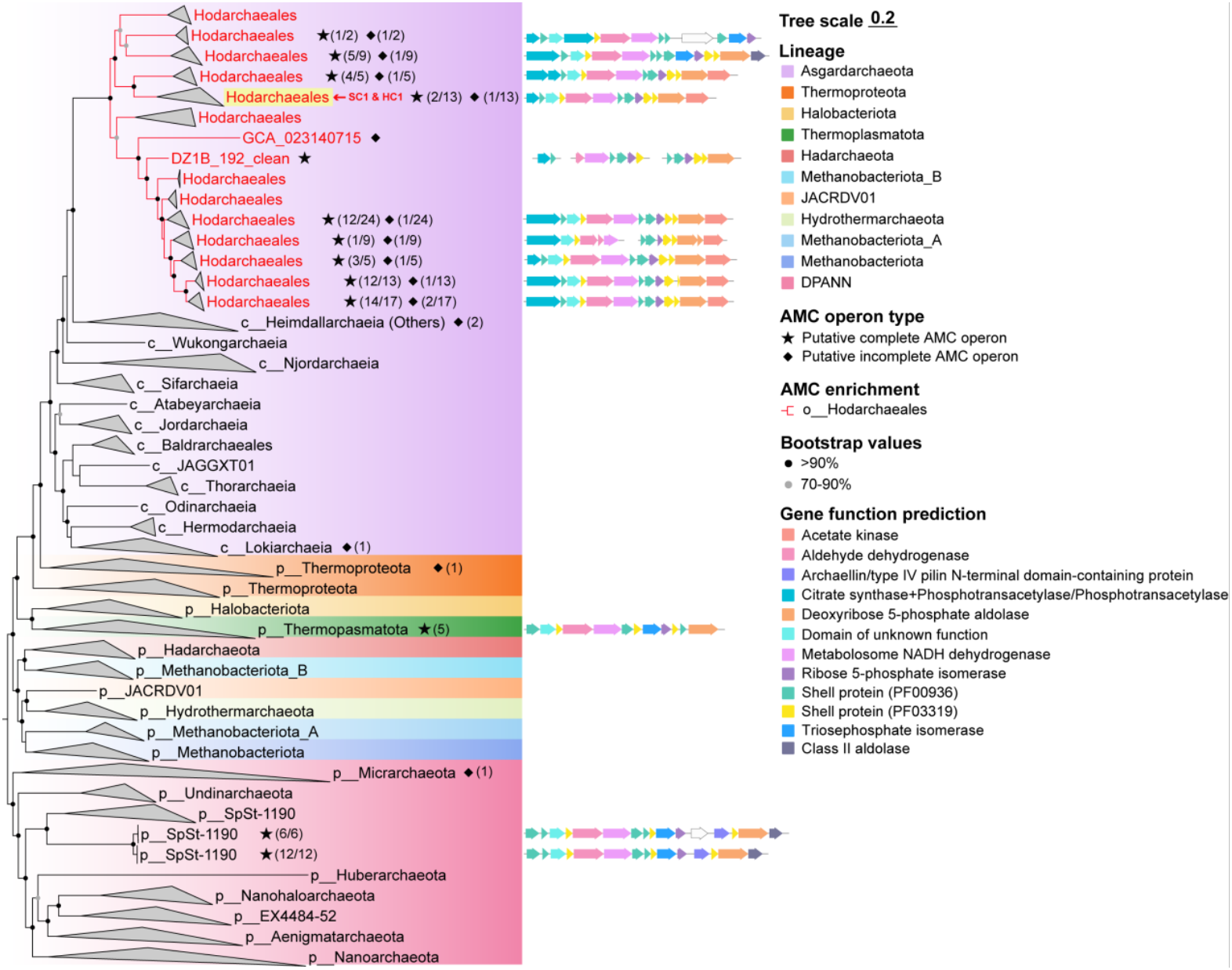
Phylogenetic distribution of the AMC operon in Archaea. A maximum-likelihood phylogenetic tree of 53 archaeal marker gene concatenation was reconstructed from 1470 representative archaeal genomes using IQ-Tree v2.3.6 (-m LG+C60+G -mwopt -B 1000 -alrt 1000) with the posterior mean site frequency (PMSF) approximation using the guide tree inferred under LG+G model. Lineages within the order Hodarchaeales are shown in red. The clade containing SC1 and HC1 (the two isolated Hodarchaeales strains) is highlighted in yellow. Symbols (star or diamond) are followed by the number of genomes containing putative complete or incomplete AMC operons (in parentheses). The denominator after the slash indicates the total number of genomes within the corresponding lineage. Putative complete AMC operons are annotated alongside the corresponding genomes (right panel).

### Horizontal transfer of Hodarchaeales AMC operons from a deep-branching lineage of uncultured Myxococcota bacteria

We performed comprehensive phylogenetic analyses to resolve the evolutionary origin of the Hodarchaeales AMC operons. The phylogenetic tree based on concatenated shell proteins and enzymes was largely consistent with the Hodarchaeales species tree topology (Fig. 2a, Supplementary Table 2), suggesting vertical inheritance rather than frequent horizontal transfer of AMC operons within the Hodarchaeales lineage. By reconciling single gene trees with the species tree, ancestral state reconstruction (ALE) analysis yielded a moderate probability of 0.29 for the presence of the AMC operon in the last Hodarchaeales common ancestor (LHoCA) node. This value approaches the minimum threshold (0.3) used to infer gene presence^24^, suggesting that LHoCA might have possessed AMC. The absence of AMC operons in certain Hodarchaeales genomes (45 of 111, 41%) is unlikely to be fully explained by metagenomic incompleteness, given that AMC-lacking genomes had comparable CheckM2 completeness scores (average 83.3%) to AMC-containing genomes (average 80.1%). Rather, this pattern probably reflects secondary losses following AMC acquisition at the LHoCA. This scenario was further supported by examining cultured isolates, where only strain SC1 retained and actively transcribed the AMC locus^14^, whereas strain HC1 lacked AMC entirely.

**Figure 2:**
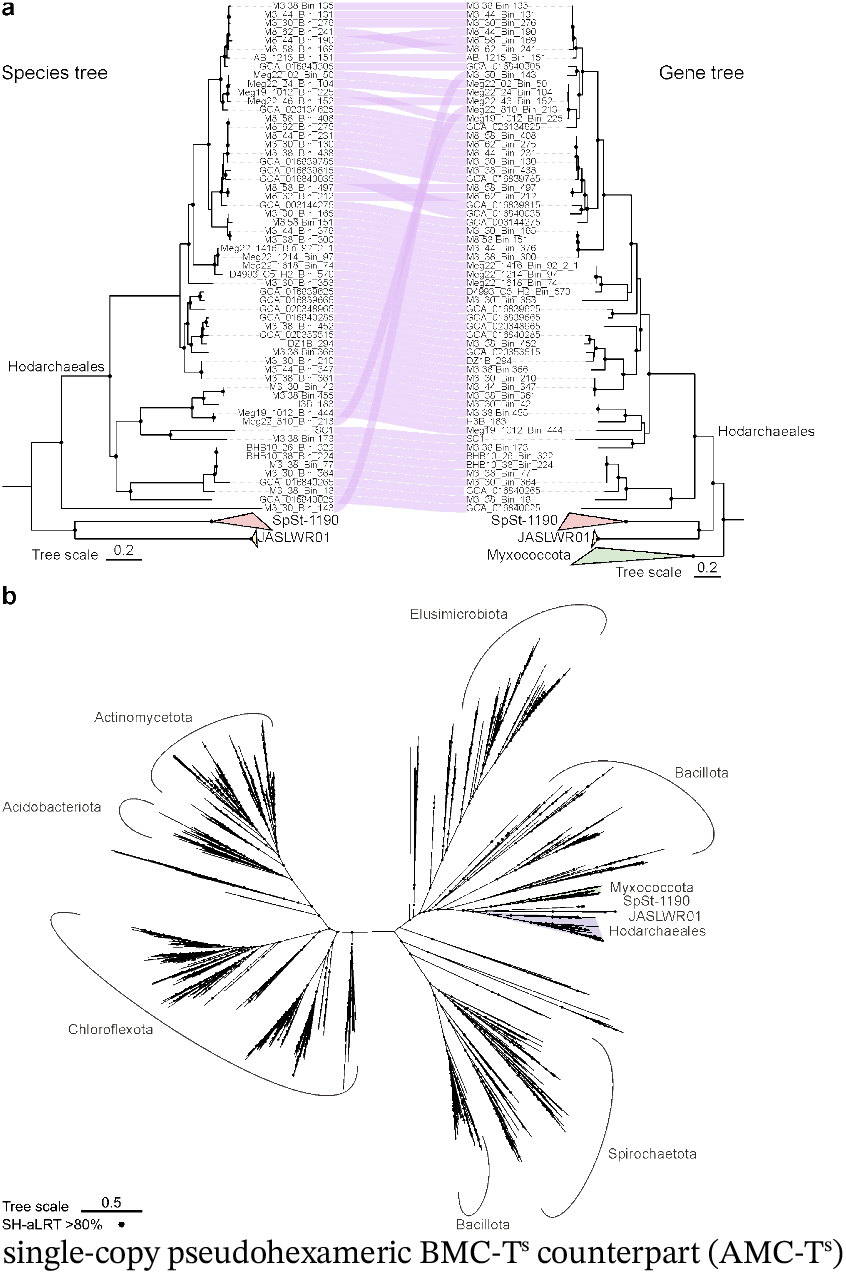
Phylogenetic analysis of archaeal microcompartments and their bacterial homologs. The maximum-likelihood phylogenetic tree was constructed using IQ-tree with LG+C60+F+G model and posterior mean site frequency (PMSF) approximation. Nodes with SH-aLRT bootstrap values greater than 95% are indicated by solid circles. **(A)** Vertical inheritance of AMC clusters within Hodarchaeales. The species tree shown in the left panel was constructed based on the multiple alignment of 53 conserved archaeal proteins following the GTDB-tk pipeline. Genomes lacking AMC clusters are not shown for clarity. The AMC gene tree in the right panel is a subtree of the tree in **(B)**. that was rooted at the Myxococcota branch. The relationships between the two trees are highlighted, with connecting lines shown in light purple. **(B)** Maximum-likelihood phylogenetic tree of prokaryotic SPU-type microcompartments built based on concatenated alignments of eight shell proteins and five enzymes (2944 amino acids in total, see details in Supplementary Table 2). AMCs and their likely Myxococcota donor are shaded according to their taxon affiliation.

Notably, in both concatenated and single-gene AMC phylogenetic trees, Hodarchaeales AMCs consistently clustered with BMCs from uncultured Myxococcota lineages (B64-G9 class and HGW-17 order), with high statistical support (Fig. 2b and Supplementary Fig. 2-9), indicating that Hodarchaeales AMCs were acquired via HGT from Myxococcota. Hodarchaeales genomes contain two types of AMCs (Fig. 3a). Because Myxococcota BMCs encompass triosephosphate isomerase (TPI) and class II aldolase characteristic of type II AMC (Supplementary Fig. 10), a parsimonious evolutionary scenario suggests that Hodarchaeales ancestors initially acquired type II AMC, which subsequently evolved into type I in most Hodarchaeales genomes during their later diversification. This evolutionary trajectory is supported by phylogenetic analyses, where type II AMCs occupy more basal positions within the Hodarchaeales clade in both concatenated and single-gene phylogenies (Fig. 2b and Supplementary Fig. 2-9). The gene transfer from Myxococcota to Hodarchaeales likely occurred early in geological time because the B64-G9 class represents a deeply branching lineage among Myxococcota (Supplementary Fig. 11). To date the HGT, we examined the GTDB relative evolutionary divergence (RED) value, which represent genetic divergence levels of certain taxa and correlate well with absolute geological time^25^. Hodarchaeales, B64-G9 class, and Myxococcota displayed RED values of 0.392, 0.449, and 0.308, corresponding to approximate origination times of 2.43 billion years ago (Gya), 2.20 Gya, and 2.76 Gya, respectively, suggesting ancient emergence of the AMC. These approximate estimates are consistent with recent molecular dating analyses, which placed Hodarchaeales or crown Heimdallarchaeia origins between 3.12-2.26 Gya^9^ or between 2.80-2.19 Gya^26^.

**Figure 3:**
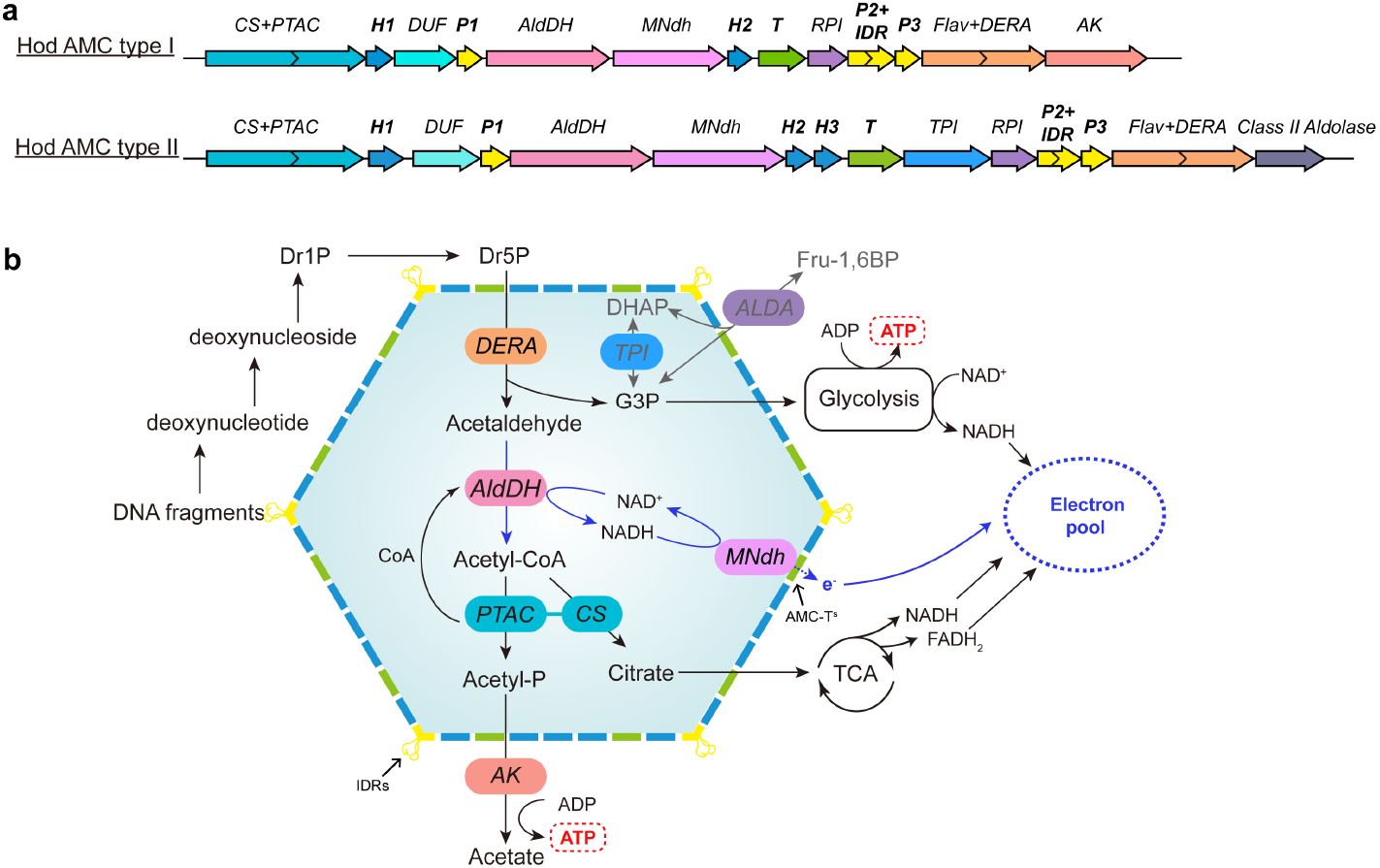
Gene organization of AMC operons in Hodarchaeales and the inferred AMC metabolism. **(A)** Two representative Hodarchaeales AMC operons consisting of genes encoding six structural components (H1, P1, H2, T^s^, P2 and P3) marked in bold along with genes encoding metabolic enzymes. **(B)** Proposed metabolic scheme for the Hodarchaeales AMC system. The DNA degradation product 2-deoxyribose-5-phosphtae (Dr5P) serves as the entry metabolite. Catabolism of Dr5P is coupled with recycling of Acetyl-CoA and reduction of NAD^+^ to NADH. Electrons released during NAD^+^ regeneration are translocated across the AMC shell via the metabolosome NADH dehydrogenase (MNdh)-AMC-T^s^ complex, contributing to the cytosolic electron pool. Other metabolites, such as glyceraldehyde 3-phosphate (G3P) and citrate, might enter additional pathways to generate NADH, further supplementing the electron pool. Abbreviations include Hod (Hodarchaeales), CS (citrate synthetase), PTAC (phosphotransacetylase), DUF (domain of unknown function), AldDH (acetaldehyde dehydrogenase), RPI (ribose 5-phosphate isomerase), Flav (flavoprotein-containing domain), DERA (deoxyribose 5-phosphate aldolase), AK (acetate kinase), TPI (triosephosphate isomerase), Dr1P (deoxyribose 1-phosphate), Acetyl-P (acetyl-phosphate), DHAP (dihydroxyacetone phosphate), Fru-1,6BP (fructose 1,6 bisphosphate), TCA (tricarboxylic acid cycle), MK (menaquinone), cyt c (cytochrome *c*).

AMCs were also detected in the JASLWR01 order (Thermoplasmatota) and SpSt-1190 lineage (DPANN), which clustered with Myxococcota BMCs and Hodarchaeales AMCs in the phylogenetic tree (Fig. 2b). The AMC operons in these archaea differ in composition from those in both Hodarchaeales and Myxococcota (Supplementary Fig. 1, 11 and Table 2). Hodarchaeales AMCs and Myxococcota BMCs both contain two copies of the shell H gene, one copy of the T^s^ gene, and three copies of the P genes. In contrast, JASLWR01 and SpSt-1190 genomes contain an additional T^dp^ gene, and JASLWR01 apparently lost one copy of shell H gene. In addition, the gene encoding phosphotransacylase (PTAC) is present in Hodarchaeales AMC and Myxococcota BMCs but absent in JASLWR01 and SpSt-1190 genomes. Given the sparse representation of AMC in these archaeal lineages, they appear unlikely to have served as intermediaries in the HGT between Myxococcota and Hodarchaeales but rather acquired these operons via independent HGT events.

### Hodarchaeales AMC are sugar-phosphate utilization metabolosomes implicated in energy conservation

The Hodarchaeales AMC components are encoded by highly conserved operons, which can be divided into two types based on the gene composition and synteny (Fig. 1, 3a and Supplementary Fig. 1). Type I AMC encompass 2 copies of the hexameric BMC-H counterpart (hereafter AMC-H), a single-copy pseudohexameric BMC-T^s^ counterpart (AMC-T^s^)and 3 copies of the pentameric BMC-P counterpart (AMC-P). Type II AMC includes an additional copy of AMC-H (AMC-H3), with sequence similarity of 42-55% with AMC-H2. Modelling the structures of the AMC proteins encoded in a representative type I Hodarchaeales AMC operon (genome accession: GCA_003144275) using AlphaFold3^27^ predicted the formation of stable homomultimers with high confidence (Supplementary Fig. 12), compatible with the self-assembly into a functional microcompartment. Structural alignments revealed high similarity between the AMC shell proteins and their BMC counterparts (mean TM-score = 0.87 computed with US-align^28^; Supplementary Fig. 13).

This pronounced structural similarity suggests that, as in the case of the BMCs, the AMC shell is formed by AMC-H and AMC-T^s^ proteins, forming hexagonal facets with central pores for metabolite transport, whereas AMC-P proteins occupy the vertices to complete the polyhedral shell assembly^29^. A distinct architectural feature was identified in 58 of the 66 (88%) AMC operons in Hodarchaeales: AMC-P2 is fused to an intrinsically disordered region (IDR) ranging from 36 to 137 amino acid residues that are positively charged due to enrichment in lysine (K) and arginine (R) (Fig. 3a and 4a, Supplementary Table 3 and 4). Given that IDRs typically mediate molecular recognition^30,31^, their fusion to an AMC-P protein suggests a distinct function at the AMC surface.

Metabolic reconstruction suggests that Hodarchaeales AMCs function as sugar phosphate utilization (SPU) metabolosomes analogous to those recently identified across diverse bacterial phyla^23,32^. The AMC operons consistently encode the enzymes of the SPU pathway including deoxyribose 5-phosphate aldolase (DERA; PF01791) and acetaldehyde dehydrogenase (AldDH; PF00171) (Fig. 1 and 3a). The entry metabolite of this pathway, 2-deoxyribose-5-phosphate (Dr5P), is typically derived from nucleic acid degradation^23,32^. Genome analysis of Hodarchaeales revealed a complete suite of enzymes involved in the generation of Dr5P from the products of nucleic acid degradation (Supplementary Table 5). This pathway begins with dephosphorylation of deoxynucleotides to deoxynucleosides, primarily catalyzed by 5-nucleotidase. The resulting deoxynucleosides are then subject to phosphorolysis, mainly with a uridine phosphorylase being the key enzyme for generating deoxyribose 1-phosphate (Dr1P) from deoxyuridine. Finally, Dr1P is isomerized to Dr5P by a phosphodeoxyribomutase, completing the pathway. Additionally, these genomes encode enzymes for upstream deoxynucleotide pool interconversion, such as nucleoside-diphosphate kinase, adenylate kinase, dCTP deaminase, and dUTP diphosphatase, which likely supply the appropriate substrates for this degradation cascade. As shown in the next section, the K/R-rich IDRs fused to the AMC-P2 proteins bind DNA. Thus, the AMC shell is predicted to participate in cytosolic DNA capture, compatible with the Hodarchaeales SPU-AMC functioning as a system to use exogenous DNA as a nutrient source.

The AMC operon composition implies that, inside the AMCs, DERA cleaves Dr5P into glyceraldehyde-3-phosphate (G3P) and acetaldehyde, and AldDH then oxidizes acetaldehyde to acetyl-CoA, with the concomitant reduction of NAD+ to NADH. A distinct feature of DERA in most Hodarchaeales AMC operons is their fusion to a flavoprotein domain (Fig. 3a and Supplementary Table 3), which is uncommon in BMC. Due to the limited permeability of the AMC shell, accumulation of NADH could inhibit AldDH activity by depleting NAD^+^, thereby trapping the reducing power within the microcompartments^33^. We identified conserved Fe-S cluster binding motifs in the metabolosome NADH dehydrogenase (MNdh) and AMC-T^s^ (CPGK) proteins encoded in Hodarchaeales AMC operons, positioned analogously to those in the structurally characterized bacterial MNdh-BMC-T^s^ complexes^34^ (Supplementary Fig. 14 and 15). Structural modeling further supports the formation of a functional MNdh-AMC-T^s^ complex (Supplementary Fig. 16), which likely facilitates the export of electrons from the intra-microcompartment NADH to the cytoplasm via a Flavin mononucleotide (FMN) cofactor and a relay of three 4Fe-4S clusters. This process would both regenerate the internal NAD^+^ pool and provide the cell with an electron pool that could be channeled into anaerobic or aerobic electron transport chains (ETCs) for ATP synthesis, highlighting the potential role of AMCs in cellular energy production.

Beyond the redox balance, the metabolosome must also regenerate the coenzyme A (CoA) moiety from its primary product, acetyl-CoA. The AMC operon encodes two enzymes, citrate synthase (CS; PF00285) and phosphotransacylase (PTAC; PF06130), both catalyzing the release of CoA through distinct routes (Fig. 3b). PTAC converts acetyl-CoA to acetyl-phosphate (Acetyl-P), potentially supporting substrate-level ATP synthesis via acetate kinase (AK; PF00871). Alternatively, CS would condense acetyl-CoA with oxaloacetate to produce citrate, which could subsequently exit the microcompartment to fuel biosynthetic pathways or the TCA cycle for NADH generation. In some lineages, the genes encoding CS and PTAC are fused into a single open reading frame (Supplementary Table 3), which is rare in BMC operons. Type II AMC operons encode additional enzymes including triosephosphate isomerase (TPI; PF00121) and class II aldolase (ALDA; PF00596) (Fig. 1, 3a and Supplementary Fig. 1). Thus, genome analysis indicates that, together, these pathways constitute a complex, integrated system for efficient utilization of DNA-derived nutrients and cellular energy production.

### Hodarchaeales AMC shell assembly and interaction with metabolic enzymes

To demonstrate *in vitro* assembly of Hodarchaeales AMC empty shells, we expressed and purified the six core shell proteins (AMC-H1, H2, P1, P2, P3 and T^s^) from a Hodarchaeales AMC operon (genome accession: GCA_003144275) individually in *Escherichia coli* (Fig. 4a), and mixed H2, T^s^ and P1 at the 3:1:3 molar ratio (see Methods). Size exclusion chromatography (SEC) purification yielded intact particles containing all three shell proteins, as confirmed by SDS-PAGE (Fig. 4a). Transmission electron microscopy (TEM) revealed multiple shell architectures ranging from ∼50 to 220 nm in diameter, with a size distribution fitting a Gaussian function N (μ=121.74 nm, σ=33.28 nm) (Fig. 4b, c and Supplementary Fig. 17). This range of sizes is closely similar to those observed for bacterial microcompartments although the AMC size was slightly larger than the reported averages for BMCs (typically, < 110 nm)^29,35–37^. The assembled shells displayed ovoid morphologies with clear boundaries and uniformly distributed pore-like structures across their shell surfaces (Fig. 4c), suggestive of the formation of intact shells. To the best of our knowledge, sugar-phosphate utilization microcompartments have not been visualized previously.

**Figure 4:**
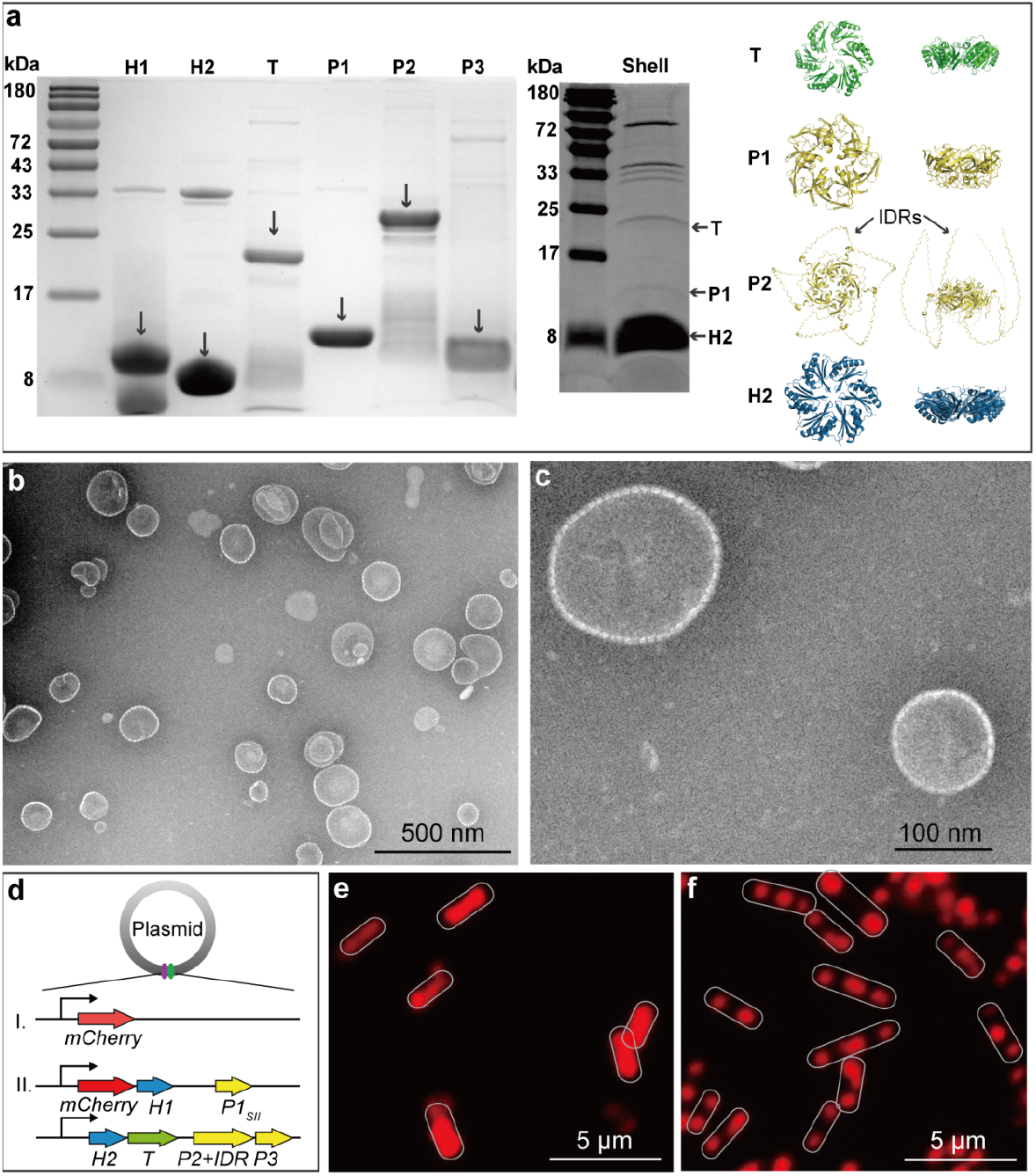
Characterization of reconstituted Hodarchaeales AMC shells. **(A)** SDS–PAGE analysis of purified individual shell proteins (H1, H2, T^s^, P1, P2, P3) and the reconstituted AMC-H2TP1 shell complex. Arrows indicate distinct protein bands. The right panel shows predicted structures of shell proteins generated with AlphaFold3. H2, T^s^ and P1 are the shell proteins constituting AMC shells. P2 is the unique shell protein with IDRs. **(B-C)** TEM images of *in vitro* assembled AMC-H2/T^s^/P1 shells. Scale bars: 500 nm (b), 100 nm (c). **(D)** Schematic of plasmid constructs designed for heterologous expression: (I) mCherry alone; (II) co-expression of mCherry with the shell proteins (H1, H2, T^s^, P1_SII_, P2, P3). **(E-F)** Fluorescence microscopy images of *E. coli* cells expressing mCherry alone (e) and co-expressing mCherry with shell proteins (f). Scale bars: 5 μm.

For *in vivo* assembly, plasmids encoding H1-P1_SII_ and H2-T^s^-P2-P3 were co-expressed in *E. coli* BL21 (DE3) (Fig. 4d). Subsequent SDS-PAGE and MS analysis confirmed the purification of intact shell particles containing all five AMC shell proteins, except AMC-H2, from the cell lysate (Supplementary Fig. 18). For intracellular visualization of shell particle assembly, we fused mCherry to the H1 protein. While mCherry alone showed diffuse cytosolic distribution, co-expression of mCherry-H1-P1_SII_ with H2-T^s^-P2-P3 yielded discrete fluorescence patches in the cells (Fig. 4e and f), compatible with shell formation *in vivo*. These results collectively demonstrate that the shell proteins encoded in Hodarchaeales AMC operons serve as building blocks for shell particle assembly.

We next investigated the interactions between core metabolic enzymes and shell proteins. Encapsulation peptides (EPs) that typically form short amphipathic α-helices and are appended as terminal extensions to the respective proteins^38^ mediate enzyme packaging into microcompartments^39–41^. AlphaFold3 structural modeling^27^ and amphipathicity analysis^38^ predicted that four of the seven core enzymes (Flav+DERA, AldDH, CS+PTAC and DUF) of the AMCs contain potential EPs located at terminal regions or as linkers between two domains of the respective enzymes (Supplementary Fig. 19). Co-expression with shell particles showed that GFP-tagged, EP-containing enzymes were selectively recruited, whereas EP-deletion variants showed no detectable interaction (Supplementary Fig. 20), confirming the essential role of AMC EPs in the packaging of these enzymes. Furthermore, we observed that two other enzymes, AK and RPI, interacted with shell proteins, despite lacking predicted EPs. Nevertheless, AK is predicted to function outside the metabolosomes^16,42,43^ presumably due to the difficulty of ATP/ adenosine diphosphate (ADP) transport across the AMC shell pore, whereas the role of metabolosome-associated RPI remains unexplored.

To investigate the function of the IDRs in the AMC shell proteins, we compared the nucleic acid-binding capacities of the full-length AMC-P2, a truncated version of AMC-P2 lacking the IDR, and the isolated IDRs, following their ex-pression *in vivo*. We found that the full-length P2 and the IDR alone but not P2 lacking the IDR bound DNA or RNA (Supplementary Fig. 21a). To further validate the nucleic acid-binding capacity of the IDRs, we treated the full-length AMC-P2 eluates with proteinase K, DNase, and RNase to degrade proteins, DNA and RNA, respectively. Subsequent agarose gel electrophoresis analysis demonstrated the presence of released DNA and RNA in the proteinase-treated eluates (Supplementary Fig. 21b), supporting the conclusion that the IDRs mediate DNA and RNA binding.

### AMC-driven energy production in Hodarchaeales

After validating the formation of Hodarchaeales AMC shells and identifying the encapsulated sugar-phosphate utilization enzymes, we sought to quantify the energetic benefits conferred by this compartmentalized metabolic system. Building on evidence that microcompartments enhance metabolic efficiency via enzyme and substrate concentration^44,45^, we hypothesized that the ability of Hodarchaeales AMCs to generate NADH and export electrons could substantially augment cellular energy production. To quantify this prediction, we developed a steady-state, one-dimensional reaction-diffusion model in spherical coordinates, under the assumption that both the cell and the AMC exhibit spherical symmetry (Fig. 5a). The model incorporated three core processes: i) conversion of DNA to Dr5P, catalyzed by uridine phosphorylase (UPP) (the rate-limiting step^46^), ii) subsequent catabolism of Dr5P to acetaldehyde catalyzed by DERA, iii) AldDH-catalyzed oxidation of acetaldehyde coupled to NADH generation. While Dr5P is generated in the cytosol, the subsequent steps are confined to the AMC lumen. A detailed description of the model and the simulations are provided in Methods and Supplementary Note 1.

**Figure 5:**
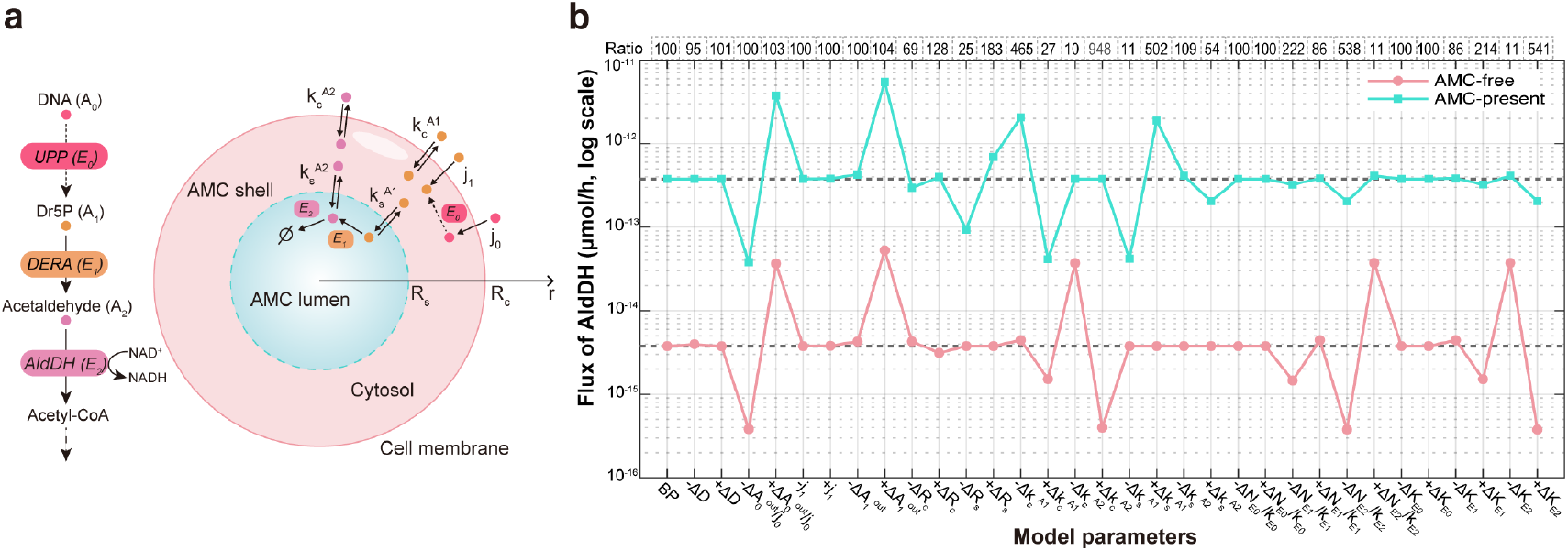
Metabolic modeling for comparing the flux through acetaldehyde dehydrogenase within the AMC and in the absence of AMC. **(A)** Schematic of the mathematical model for metabolic pathways associated with the AMC. Extracellular DNA (A_0_) enters the cell with active transport (rate j_0_) and is converted to 2-deoxyribose-5-phosphtae (Dr5P, A_1_) through a rate-limiting step catalyzed by uridine phosphorylase (UPP, E_0_) in the cytosol. Dr5P is transported across the cell membrane via active transport (rate j_1_) and channel-mediated facilitated diffusion (rate k_c_^A1^), and across the AMC shell via permeability-dependent diffusion (rate k_s_^A1^). Within the AMC, Dr5P is converted to acetaldehyde (A_2_) by deoxyribose 5-phosphate aldolase (DERA, E_1_), which is then converted to acetyl-CoA with concomitant NAHD generation via acetaldehyde dehydrogenase (AldDH, E_2_). Acetaldehyde diffuses across the AMC shell and cell membrane with permeability rates k_s_^A2^ and k_c_^A2^, respectively. R_c_ and R_s_ denote the cell and AMC radius, respectively. **(B)** Comparison of metabolic flux through AldDH within the AMC and in the absence of AMC. Each parameter was varied up to 10-fold from the estimated value (Baseline Parameter, BP), except for the cell and AMC radius (see Supplementary Note 1).

To assess the potential role of the AMC in energy conservation, we compared the metabolic flux through AldDH within the AMC to that in the cytosol in the absence of this microcompartment. With the estimated model parameters (Supplementary Note 1), AMC-containing systems showed an approximately 100-fold increase in AldDH-mediated flux and the corresponding NADH generation (Fig. 5b). To account for potential variations in parameter values under different environmental conditions, we performed simulations with each parameter perturbed 10-fold, either upward or downward. In these simulations, the AMC-containing systems consistently outperformed their cytosolic counterparts, with flux enhancements ranging from 10-fold up to nearly 1000-fold across broad parameter ranges, demonstrating the robustness of the potential energy production boost provided by compartmentalization. The AldDH flux within AMC showed lower sensitivity to the kinetic parameters, reflecting the enhancement of the enzymatic efficiency by the microcompartments. Furthermore, the lower cytosolic acetaldehyde concentration in the model including AMC (1% of that in the model without AMC) suggested that AMCs also protect the cells by sequestering the toxic acetaldehyde.

During the period of fluctuating oxygen levels, the energy surplus provided by the AMC supported diverse ESP-mediated functions and cellular morphological changes under two alternative scenarios. Anoxic scenario: ESP function was sustained under anoxic conditions, powered by an anaerobic electron transport chain (ETC) that was continuously supplied with electrons form AMCs. In this context, the UOX/COX system would have functioned primarily for antioxidant defense when the cell encountered transient oxygen. Microoxic scenario: In microoxic environments, ESP activity was driven by a highly efficient aerobic ETC that integrated electrons from the AMC for maximal energy yield. Upon oxygen depletion, the host would revert to the AMC-fueled anaerobic ETC to survive the environmental shift. IV. Engulfment of the alphaproteobacterial ancestor of the mitochondrion. Microoxic conditions facilitated the coexistence of the Hodarchaeales ancestor and the mitochondrial ancestor, and the high energy yield provided by the AMCs supported engulfment of the alphaproteobacterium. V. Establishment of endosymbiosis. Following the engulfment of the mitochondrial ancestor, a stable endosymbiotic relationship was established over an extended evolutionary period, culminating in full integration of the mitochondrion with the concomitant loss of the AMC.

## Discussion

Although a variety of eukaryogenesis models centered around the proto-mitochondrial symbiosis have been proposed ^5–7,47–49^, a key unresolved question is how the archaeal host (or FECA) surmounted the bioenergetic barrier to fuel the evolution of eukaryotic complexity, such as evolution of the molecular machinery for membrane remodelling and cytoskeleton which are of apparent Asgard origin^2–4^. FECA might have inhabited both anoxic and microoxic niches^9,13,14^, cohabitation with the alphaproteobacterial ancestor of the mitochondrion likely occurring in the latter. Among several proposed mechanisms, phagocytosis is usually considered most plausible for the acquisition of the proto-mitochondrial ^7,13,50^. Phagocytosis is generally considered to be a eukaryotic staple, but recent observations on large bacteria^51^ and particularly a phagocytic Planctomycetotal bacterium^52^ indicate that prokaryotes, in principle, can be phagocytes. Together with the complementary evidence of innate complexity in Asgard archaea, extensive ancient gene duplications^53^, and the apparent dominant contribution of Asgard archaea to the origin of eukaryotic genes^54^, these findings appear to be best compatible with mito-intermediate models of eukaryogenesis. Such models posit that the archaeal ancestors of eukaryotes evolved some cellular complexity prior to and independent of the acquisition of the proto-mitochondrial symbiont, setting the stage for productive endosymbiosis^13,47–49^. Phagocytosis, however, is bioenergetically demanding^5,10,55–57^.

An important question, then, is what metabolic devices in the archaeal host could have provided the necessary bioenergetic foundation for the transition to the eukaryotic cellular complexity. Here, we report the discovery of complex AMCs in Hodarchaeales, the apparent descendants of the FECA, which provide a potential solution to this bioenergetic problem. Unlike the previously reported protein-bounded nanocompartments (e.g., encapsulins) in Asgard^14^ and other archaea^58,59^, the Hodarchaeales AMC components are encoded in a conserved operon comprising self-assembling shell proteins, a set of metabolic enzymes, and electron transport chain subunits. Such microcompartments endowed with multi-enzyme metabolic complexity have been previously observed in bacteria but, to our knowledge, not in archaea. The AMCs most likely function to enhance energy production from scavenged DNA. Mathematical modelling showed that AMCs can boost NADH yields about 100-fold compared to non-compartmentalized pathways (Fig. 5b), primarily, through substrate channelling and enzyme colocalization. This efficiency is apparently sustained by an internal electron transfer channel, the MNdh-AMC-T^s^ complex. This complex potentially resolves a fundamental constraint of AMCs, whose narrow shell pores (≤5 Å in diameter)^19,60^ exclude NAD^+^/NADH diffusion, by exporting electrons derived from compartmentalized NADH oxidation directly to the cytosol^34^. This process maintains a separated, sequestered, regeneratable luminal NAD^+^/NADH pool^34^, while providing the cell with a substantial flux of reducing power. These electrons might then fuel high energy production by reducing cytosolic acceptors, such as ferredoxins or O_2_^61,62^, thereby providing another possible mechanism for oxygen detoxification in Hodarchaeales^14,15^. Intriguingly, TEM revealed ferritin-like structures in strain HC1^14^, suggesting that AMCs might function synergistically with iron storage systems to maintain redox homeostasis, a potential adaptation to fluctuating oxygen conditions. Although *in vivo* AMC structures have not been visually observed in Hodarchaeales strain SC1, transcriptomic data indicates active expression of AMC operons, with AMC-H2 among the top 3% most highly expressed proteins^14^.

Perhaps, unexpectedly, genomic comparisons reveal substantial differences between the energy conversion and antioxidant systems in the two available Hodarchaeales isolates. Strain SC1 possesses the AMC gene cluster but lacks globin, whereas strain HC1 encodes globin and ferritin but lacks the AMC cluster, apparently having lost it during evolution. Thus, different representatives of Hodarchaeales seem to employ distinct strategies to manage redox homeostasis and energy generation under fluctuating oxygen conditions.

The architecture of the AMC reveals a potent bioenergetic engine apparently adapted to an ecological niche with ample environmental DNA supply and fluctuating oxygen levels. A notable innovation of Hodarchaeales AMCs is the fusion of IDRs to the vertex-forming AMC-P pentamers, a feature not observed in BMCs. Our data show that these K/R-rich IDRs confer the DNA-binding capacity to the AMC shells (Supplementary Fig. 21), enabling the AMC to function as a dedicated apparatus for capturing and processing environmental DNA. This DNA-centric metabolism is well-suited for a lifestyle in sediments, where extracellular DNA persists as a stable and abundant nutrient supply^63,64^. Indeed, all known Hodarchaeales lack *de novo* purine biosynthesis pathways (Supplementary Table 7) and thus should depend on exogenous nucleotides. The IDR-mediated DNA recruitment establishes a direct mechanistic link between cytosolic scavenging and compartmentalized catabolism of exogenous DNA. This system likely functions synergistically with DNA uptake mechanisms that should be highly efficient given the high level of HGT documented in Asgard archaea^65,66^.

The LHoCA might have originated close to the Great Oxygenation Event^9^. To cope with the increasing oxygen concentration, Hodarchaeales ancestors apparently evolved multiple adaptive strategies including oxygen-reducing enzymes (e.g., COX), which function in antioxidant defense and/or aerobic respiration^13^, and AMCs, which are predicted to promote energy generation under both anaerobic and aerobic conditions to adapt to fluctuating oxygen levels. Hodarchaeales ancestors might have acquired COX and SPU-AMC within the same, short time interval around the Great Oxygenation (Supplementary Fig. 22), from aerobic archaea and Myxococcota, respectively. Recent phylogenomic analysis suggests that Myxococcota-derived genes contributed to LECA at levels comparable to those from Alphaproteobacteria, particularly enriching pathways involved in nucleotide biosynthesis, nucleoside-sugar metabolism, and steroid precursor synthesis^54^, partially overlapping with AMC functions and suggestive of a deep evolutionary connection. Such an association between the likely archaeal ancestors of eukaryotes and Myxococcota are compatible with the syntrophy model of eukaryogenesis in which archaeal-myxococcal symbiosis is postulated to precede the capture of the proto-mitochondrion^42,44^. The AMC system likely increased fitness by enabling the exploitation of environmental DNA as both a carbon source and an electron sink, which could be pivotal during the Proterozoic era when fluctuating oxygen levels^67^ favoured metabolically flexible organisms—traits later co-opted during eukaryogenesis.

Based on our findings, we propose a hypothetical model for the possible role of AMCs in eukaryogenesis (Fig. 6). Our ancestral reconstruction suggests that the LHoCA co-acquired AMCs and UOX/COX terminal oxidases (Supplementary Fig. 22-23) through horizontal gene transfer from Myxococcota and aerobic archaea, respectively. The capture of these genes likely provided a substantial metabolic advantage by compartmentalizing sugar-phosphate metabolism, enabling Hodarchaeales to thrive during the prolonged anoxic-to-microoxic transition period preceding the Great Oxygenation. Within this framework, we propose two alternative scenarios for the establishment of the symbiosis with the mitochondrial ancestor, depending on the prevailing oxygen availability at the time.

**Figure 6:**
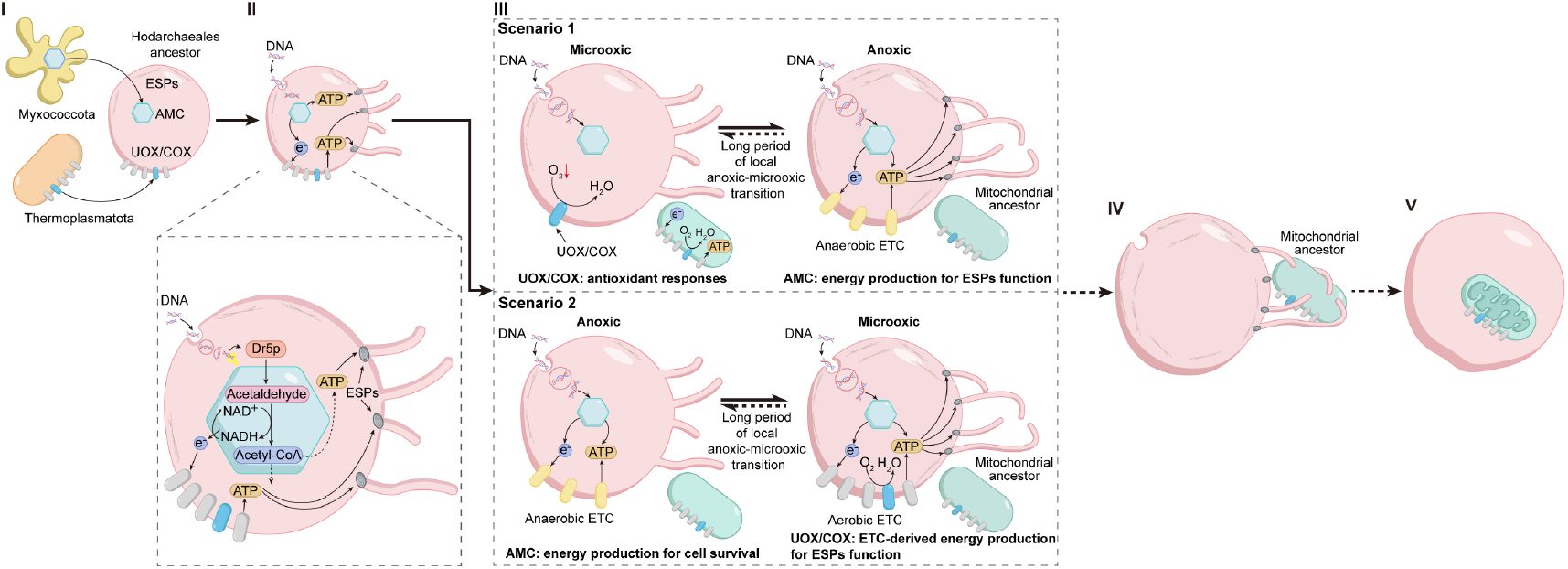
AMC-energy driven model of eukaryogenesis. The AMC (archaeal microcompartment)-energy driven model consists of five stages, from the Last Hodarchaeales Common Ancestor (LHoCA) mitochondrial endosymbiosis and the LECA. I. Acquisition of essential metabolic systems. The LHoCA acquired two key systems via HGT: AMCs from Myxococcota, possibly, through a syntrophic association, and a UOX (ubiquinol oxidases)/COX (cytochrome c oxidase) complex from Thermoplasmatota. II. Fueling evolution of cellular complexity. The AMCs provided a substantial bioenergetic surplus which was a prerequisite for powering the evolution of complex cellular machinery involved membrane remodeling and cytoskeleton functions and largely consisting of Eukaryotic Signature Proteins (ESPs). III. Adaptation to anoxic-microoxic transition conditions.

The first scenario posits that the pre-mitochondrial endosymbiosis occurred under strictly anoxic conditions. In the anoxic environment, the LHoCA cell would leverage the AMCs for scavenging external DNA to generate electrons, which would be funnelled into an anaerobic ETC for energy production. Upon transitioning to a microoxic environment, the LHoCA would utilize the UOX/COX for antioxidant defense, detoxifying reactive oxygen by reducing it to water. The terminal oxidases would be essential for survival but might not have played a major role in energy generation. In this scenario, AMCs would have continued to account for a large proportion of cellular energy generation.

Alternatively, the pre-mitochondrial endosymbiosis might have occurred in a microoxic environment. In this case, the key evolutionary leap was the host’s adaptation to these changing conditions. The host cell would repurpose its UOX/COX complex, shifting its function from simple oxygen detoxification to serving as the terminal oxidase in the aerobic ETC. While the AMC’s core function of DNA breakdown would have persisted, the produced electrons would be channelled into the highly efficient aerobic pathway, fuelling more complex cellular innovations. This system would also provide crucial flexibility in the face of the fluctuating oxygen concentration; once the environmental oxygen was depleted, the LHoCA would switch to its anaerobic ETC, relying on the AMC to persist through the environmental shift. Under both scenarios, the integration of AMC-derived electrons into either the anaerobic or the aerobic ETC generated an essential bioenergetic surplus, which may have been important for powering eukaryotic signature proteins (ESPs) involved in membrane remodelling and cytoskeleton formation, and thus setting the stage for subsequent eukaryogenesis.

## Conclusion

Taken together, our results suggest that Hodarchaeales AMCs represent a pivotal innovation providing the bioenergetic efficiency that enabled critical pre-adaptations for eukaryogenesis. Through the horizontal acquisition of a Myx-ococcota-derived microcompartment system, ancestral Hodarchaeales gained: (i) a DNA-capturing apparatus for nutrient exploitation in nucleic acid-rich environments, (ii) enzymatic compartmentalization enabling high efficiency of NADH production and regeneration, and (iii) an enhanced energy production via electron export from the AMC to the cytosol. This metabolic module would enable the adaptation of the anaerobic ancestors of Hodarchaeales to Proterozoic oxygen fluctuations and the emergence of biochemical and subcellular complexity via enhanced energy production. Compartmentalization derived through HGT from bacteria might have provided the archaeal host with the bioenergetic capacity, redox stability, and metabolic flexibility necessary to support the acquisition of mitochondria.

## Acknowledgements

We thank Masaru K. Nobu for providing early access to the HC1 and SC1 genomes. The work was supported by the National Natural Science Foundation of China (No. 32370055, No. 32393970, No. 32225003, No. 32200099 & No. 92051102), the General Program supported by Shenzhen Natural Science Foundation in Basic Research Fund (No. JCYJ20230808105711023), the Guangdong Basic and Applied Basic Research (2025A1515012817), the Project of Department of Education of Guangdong Province (2025KCXTD039), Shenzhen University 2035 Program for Excellent Research (2022B002) and the Synthetic Biology Research Center of Shenzhen University. This work was also supported by the Moore-Simons Project on the Origin of the Eukaryotic Cell, Simons and Moore Foundation 73592LPI to B.J.B. (https://doi.org/10.46714/735925LPI) and Simons Foundation Investigator in Aquatic Microbial Ecology Award (LI-SIAME-00002001). E.V.K. is supported by the Intramural Research Program of the National Institutes of Health of the USA. This paper was typeset with the bioRxiv word template by @Chrelli: www.github.com/chrelli/bioRxiv-word-template.

## Author contributions

Y.L., H.D. and M.L. conceptualized and designed the study. H.D. performed bioinformatic mining, structural modelling and mathematical modelling. A.X. and H.L. performed molecular cloning, protein characterization and microscopy analysis. X.F. and W.H. performed phylogenetic analysis. Y.L., H.D., A.X., X.F., and M.L. wrote the manuscript with help from all authors.

## Materials and Methods

### Bioinformatic mining for microcompartment signature in archaeal genomes

We analyzed 15,821 archaeal genomes from several sources: GlobDB (release 220), the Global Ocean Microbiome Genome Catalogue (GOMC; https://db.cngb.org/maya/datasets/MDB0000002) and additional Asgard genomes from recent studies^9,13,14^. Genomic data were standardized using anvi’o (development version 8) according to the protocol of GlobDB (https://globdb.org/methods) to ensure uniformity in downstream analyses. Specifically, genes were predicted with Prodigal (v2.6.3) implemented within the anvi’o platform using default parameters. Functional annotation of protein-coding sequences was performed with Pfam (v37.0) to identify conserved protein domains. To identify putative archaeal homologs of bacterial microcompartment (BMC)-related operons, we used hmm profiles to scan for hallmark BMC Pfam domains (PF00936 and PF03319). For each identified target locus, we extracted flanking genomic regions spanning the same strand open reading frames (ORFs) within 15 genes upstream or downstream of each target locus. Co-directionality and synteny were enforced to reconstruct operon architectures. Redundant genomic segments were collapsed to generate non-overlapping loci. Orthologous groups of BMC-associated loci were inferred using OrthoFinder (v3.0.1b1) with MMseqs2 for sequence similarity searches and Markov clustering (inflation factor = 3). For functional annotation of orthologous groups, medoid sequences—representing the centroid of multiple sequence alignments generated with MAFFT-linsi (v7.525)—were queried against the NCBI CD-search server (conserved domain database), Pfam, UniProt, and a custom database of BMC-associated HMM profiles (https://www.kerfeldlab.org/bmc-locus-hmms.html). This multi-layered annotation framework integrated domain architecture, sequence homology, and phylogenetic context to resolve evolutionary relationships and functional potential of archaeal BMC-like operons. The metabolic pathways in AMC-containing Hodarchaeales genomes upstream or downstream associated with AMC, such as DNA degradation to Dr5P and redox enzymes involved in the electron transport chain, were annotated using kofam (version 2024-10-07) with a coverage threshold of 0.7 and an E-value cutoff of 0.001. Identified loci were classified as putative complete or incomplete SPU-AMC systems based on the presence and completeness of shell proteins (P, H, H^p^, T^s^, T^sp^, and T^dp^) and core enzymes (OG0000002/AldDH and OG0000006/DERA; see Supplementary Table 2). Putative complete SPU-AMC loci were defined by the simultaneous presence of complete shell proteins (both P and H components) and complete core enzymes (both OG0000002/AldDH and OG0000006/DERA). All other scenarios, including loci with partial shell proteins or missing core enzymes, were classified as incomplete SPU-AMC loci. Incomplete SPU-AMC loci, which may result from the incomplete metagenome-assembled genomes, were excluded from the main analysis but are documented in Supplementary Table 2.

### Phylogenetic analysis

For the construction of the archaeal species tree, we sampled 1160 representative genomes from the genome taxonomy database release 220^68^ by randomly sampling 5 taxa from each family with best genome quality (completeness-5*contamination). To better represent diversity of this lineage, we expanded the taxonomic sampling in Heimdallarchaeia based on the results and genomes from a recent study^13^. We also included additional archaeal genomes encoding microcompartment operons in the species tree reconstruction, bringing the total number of archaeal genomes in the species tree to 1473. Using “identify” function in gtdbtk v2.4.0^69^, we extracted marker protein homologs assigned to the 53 archaeal marker genes. Alignments were generated using mafft-linsi v7.453^70^. Poorly aligned regions were trimmed with BMGE v1.12 (-m BLOSUM30 -h 0.55)^71^. The species tree of Archaea was inferred using IQ-Tree v2.3.6^72^ (-m LG+C60+G -mwopt -B 1000 -alrt 1000) with posterior mean site frequency (PMSF)^73^ approximation using a guide tree inferred under LG+G model and was illustrated using TVBOT v2.6.1^74^. The species tree phylogenetic analysis was performed for Myxococcota similarly based on GTDB-Tk alignment of 120 conserved bacterial proteins and IQ-Tree (-m LG+G).

For gene tree phylogenetic analysis of SPU microcompartment cluster, the five enzymes (PF00171, PF01512, PF01791, PF02502, and PF06130) and eight types of shell proteins (T^dp^, T^s^, two types of H, and four types of P)^23^ each were aligned using MAFFT-linsi v7.525^70^ and trimmed using ClipKIT v2.4.1^75^ with the ‘kpic-gappy’ parameter. Single gene trees were inferred using IQ-TREE v2.4.0^72^ (-m LG+I+G). These alignments were further merged into a supermatrix of 2944 amino acids. The concatenated gene tree was constructed using IQ-TREE v2.4.0 (-m LG+C60+I+G -alrt 1000) with PMSF approximation using a guide tree inferred under LG+I+G model. To compare the topology similarity between species tree and AMC cluster within Hodarchaeales, genomes containing AMC were extracted using the nw_prune function from Newick Utilities^76^ and compared to AMC topology using iTOL v6^77^.

### Gene gain and loss reconstruction

To minimize the potential impact of genome incompleteness on ancestral reconstruction, Heimdallarchaeia genome quality was evaluated using CheckM^78^ v1.0.2 and only those with completeness >80% were retained. The species tree, AMC gene trees and COX gene trees of high-quality genomes were extracted using nw_prune. Reconciliations between the species tree and gene trees were performed using ALEobserve (v.1.0) and ALEml_undated^79^.

The stem lengths (sl) of AMC and COX subunits at the LHoCA node were calculated following a previous study^80^. Briefly, the stem length was determined as the raw stem length of LHoCA divided by the median distance from terminal nodes to LHoCA. The stem length values of each AMC gene and COX subunit were then statistically compared using the Wilcoxon sum-rank test.

The relative evolutionary divergence (RED) in the GTDB database has been reported to correlate well with absolute geological time^25^. To provide approximate estimates of Hodarchaeales AMC origination times, we extracted RED values for Hodarchaeales, class B64-G9, and Myxococcota from GTDB R226 and performed linear conversion to absolute geological time according to previous molecular dating analysis, where RED of 0 corresponds to 4.0 Gya and RED of 1 corresponds to present day^25^.

### IDRs and EPs prediction

All amino acid sequences of Hodarchaeales AMC-P were extracted and subjected to IDR prediction using IUPred3^81,82^. Residues with a score above 0.5 were classified as IDRs. For EP prediction, the 3D structures of the cargo proteins were modeled using AlphaFold3. Amino acid sequences corresponding to α-helices with pLDDT < 90, located at either the N-terminus, C-terminus, or between two domains of a protein, were identified as putative EPs and subsequently extracted^38^. The amphipathicity of each α-helical segment was evaluated using the amphihelix_0.2.2_py3.3.py script (https://www.kerfeldlab.org/amphihelix-scripts-and-files.html) to further support EP prediction, as EPs typically exhibit amphipathicity between 0.3 to 0.5.

### Construction of plasmids carrying the AMC operon

Codon-optimized genes AMC-H1 (CEE45_05300) and AMC-H2 (CEE45_05325) were synthesized by General Biol (China) and cloned into pET-11b using BamHI and NdeI sites. AMC-T^s^ (CEE45_05330) was synthesized and cloned into pET-30a with an N-terminal 6xHis-tag. Genes AMC-P1 (CEE45_05310), AMC-P2 (CEE45_05340), and AMC-P3 (CEE45_05345) were also synthesized and individually cloned into pET-30a, each fused to a C-terminal Strep-tag II (WSHPQFEK). All AMC shell genes originated from a representative Hodarchaeales genome (Accession number: GCA_003144275).

To construct the shell protein co-expression system, AMC-H1 and AMC-P1 gene sequences were inserted downstream of the two T7 promoters in the pRSFDuet-1 plasmid using double restriction enzyme digestion. Similarly, the AMC-H2-T^s^ and AMC-P2-P3 gene sequences were cloned into the two T7 promoter sites of the pCDFDuet-1 plasmid. Both plasmids were co-transformed into *E. coli* BL21(DE3).

For the shell-enzyme co-expression system, genes encoding the enzyme core components were codon-optimized and cloned into the pET-16b vector with C-terminal GFP fusions. This plasmid was electroporated into the strain already harboring the shell protein co-expression system, resulting in a triple-plasmid co-expression system.

### Protein expression and purification

All plasmids were transformed in *E. coli* BL21(DE3) strain (Takara Bio, Japan). Competent cells were mixed with 1 μL plasmid DNA and incubated on ice for 30 min. The mixture was heat-shocked at 42°C for 60 sec, then placed on ice for 3-5 min. Subsequently, 800 μL of sterile LB broth was added, and the culture was shaken at 37°C for 1 h. A total of 150 μL of recovered cells were spread onto LB agar plates containing appropriate antibiotics and incubated overnight at 37°C.

Transformed strains were cultured in 800 mL of LB broth at 37°C with shaking at 220 rpm and were induced at an OD_600_ of 0.6 by adding 400 μL of 1 mol/mL IPTG (Isopropyl β-D-1-thiogalactopyranoside). Cultures were further incubated at 15°C with shaking at 160 rpm for 16-18 hours. Cells were harvested by centrifugation at 5,000 rpm for 20 min at 4°C, and the resulting pellets were stored at -80°C.

The purification of AMC-H were adapted from Range *et al*.^83^. Frozen cell pellets were resuspended in 30 mL of Lysis Buffer (50 mM Tris−HCl pH 8, 100 mM NaCl, 10 mM MgCl_2_) supplemented with 100 μL of 2 mg/mL DNase I, and 25 μL of 50 mg/mL lysozyme (Yeasen, China). Cells were lysed by sonication on ice at 180 W power until complete clarification. The lysate was then treated with 900 μL Triton X-100 (3% v/v final concentration) and incubated at room temperature for 20 min with orbital shaking. Insoluble material was separated by centrifugation for 20 min at 15,000 rpm (Eppendorf Centrifuge 5424R). The supernatant was discarded, and the pellet was resuspended in 30 mL of Wash Buffer (50 mM Tris−HCl pH 8, 100 mM NaCl, 10 mM MgCl_2_, 3% v/v Triton X100), avoiding disturbing the brown cellular debris. The suspension was aliquoted into 2 mL centrifuge tubes and subjected to iterative wash cycles with Wash Buffer until a white pellet was obtained. The pellet was washed with 2 mL Lysis Buffer to remove residual Triton X-100. After decanting, the pellet was resuspended in Urea Buffer (50 mM Tris−HCl pH 8, 150 mM NaCl, 1 M urea) and incubated overnight at room temperature with gentle shaking. The suspension was centrifuged at 13,500 × g for 10 min, after which the supernatant was carefully transferred to a fresh tube while avoiding pellet disruption.

For AMC-T^s^ purification, frozen cell pellets were resuspended in 25 mL Buffer A (20 mM Tris-HCl pH 7.8, 300 mM NaCl) with the addition of 25 μL of 50 mg/mL lysozyme (Yeasen, China). Cells were lysed by sonication on ice at 180 W power, and debris was removed by centrifugation at 14,000 rpm for 30 minutes at 4°C. The clarified lysate was located onto a 1 ml HisTrap column (Cytiva, USA), equilibrated with 4% Buffer B (20 mM Tris-HCl pH 7.8, 300 mM NaCl and 500 mM imidazole). The column was washed with ten-column volumes of 10% Buffer B until A_280_ reached baseline. Proteins were then eluted with ten-column volumes of Buffer B with a gradient of 10-100% and buffer-exchanged into Assembly Buffer (50 mM Tris−HCl pH 8, 150 mM NaCl) using 10 kDa MWCO spin filters (Vivaspin, Germany).

For purification of AMC-P (P1_SII_, P2_SII_ and P3_SII_, clarified lysates were applied to a 1 ml StrepTrap XT column (Cytiva, USA) equilibrated with Buffer C (100 mM Tris-HCl pH 8.0, 150 mM NaCl, 1mM EDTA) and washed with ten-column volumes of Buffer C. Proteins were eluted with ten-column volumes of Buffer D (100 mM Tris-HCl pH 8.0, 150 mM NaCl, 1mM EDTA, 50 mM biotin) according to the manufacturer’s instructions. Eluates were concentrated and buffer-exchanged into Assembly Buffer using 10 kDa MWCO spin filters (Vivaspin, Germany).

The shell-enzyme co-expression system was purified as previously described. Briefly, lysates from co-transformed strains were clarified and loaded onto a 5 mL StrepTrap column (Cytiva, USA). After extensive washing, bound complexes were eluted with biotin-containing buffer. Eluates were concentrated using 10 kDa MWCO spin filters (Vivaspin, Germany).

### In Vitro assembly and purification

In our screening for optimal *in vitro* assembly conditions, we evaluated two distinct combinations of the H, T, and P proteins, including H1T^s^P1 and H2T^s^P1. The mixture of H2, T^s^, and P1 at a stoichiometric ratio of 3:1:3 yielded more efficient assembly. Specifically, AMC-T^s^ and AMC-P1 were mixed at a 1:3 molar ratio in ice-cold Assembly Buffer supplemented with 10% glycerol and incubated for 30 sec. AMC-H2 was then added at a molar ratio of 3:1 relative to AMC-T^s^, resulting in a final stoichiometry of H2:T^s^:P1 = 3:1:3, followed by 1 min of mixing. The final mixture was incubated at 4 °C for 12–16 hours. After assembly, the solution was filtered through a 0.22 μm PVDF membrane and loaded onto a Superdex 200 Increase 10/300 GL column pre-equilibrated with Assembly Buffer (50 mM Tris-HCl, pH 8.0, 150 mM NaCl, 10% glycerol). Size exclusion chromatography was performed at 0.5 mL/min, and peak fractions corresponding to assembled shells were collected based on A_280_. The composition of fractions was verified by SDS-PAGE.

### SDS-PAGE analysis

Proteins were separated via sodium dodecyl sulfate polyacrylamide gel electrophoresis (SDS-PAGE) on 12.5% polyacrylamide gels and stained with Coomassie Blue Staining Solution.

### Western blot analysis

After electrophoresis, the protein-containing gel was transferred onto a 0.45 μm PVDF membrane. The membrane was first briefly soaked in 100% methanol for 20 sec and then immersed in transfer buffer. All transfer components (gel, PVDF membrane, filter papers, and sponge pads) were presoaked in transfer buffer for 10 min prior to electrotransfer. Protein transfer was performed at 400 mA for 35 min under ice-cooled conditions using rapid transfer buffer (Epizyme, China). After the transfer, the membrane was immediately blocked overnight at 4°C in TBST containing 10% non-fat dry milk to prevent non-specific binding. After three 5-min TBST washes to remove blocking solution, the membrane was probed with anti-GFP monoclonal antibody (HUABIO) for 1 hour at room temperature. Unbound antibody was removed by 3–5 additional TBST washes (5 min each). The membrane was then incubated with HRP-conjugated goat anti-mouse IgG (H+L) for 1 hour at room temperature, followed by a final series of TBST washes. Target proteins were detected using ECL substrate and visualized with a chemiluminescence imaging system.

### Transmission Electron Microscopy (TEM) Analysis

A 10 μL sample was applied to a carbon-coated copper grid (200 mesh) for 1 min. Excess liquid was wicked away with filter paper and stained with 10 μL of uranyl acetate for 1 min. The stain was blotted off, and the grid was air-dried at room temperature. Images were acquired using an 80 kV Hitachi HT7800 TEM and processed using ImageJ software^84^.

### Structural modelling

The structure for proteins were predicted using Alphafold3 (https://golgi.sandbox.google.com/)^27^. Protein structures were visualized with PyMOL (The PyMOL Molecular Graphics System, version 2.6.0 Schrödinger, LLC)^85^.

### Confocal Microscopy

Imaging was performed on a Nikon A1 HD25 confocal microscope with a 100× oil-immersion objective. Excitation wavelengths of 488 nm and/or 561 nm were applied. Images were acquired at a resolution of 512×512 using NIS software. The pinhole was set to 1.2 AU. All colocalization imaging results were analyzed using ImageJ^84^.

### Mathematical modelling of the metabolic flux within AMC

To simulate metabolic flux, we constructed a one-dimensional, steady-state reaction-diffusion model in spherical coordinates. The model was designed to compare metabolic outcomes under two distinct scenarios: (i) an AMC-present case, where part of the key enzymatic reactions was compartmentalized within the AMC, and (ii) an AMC-absent case, where all reactions were assumed to occur in the cytosol.

The model incorporated three core reactions converting DNA to Acetyl-CoA: (i) the DNA-to-Dr5P conversion catalyzed by uridine phosphorylase (UPP, denoted as *E*_0_), which was assumed to occur exclusively in the cytosol in both scenarios, (ii) the conversion of Dr5P to acetaldehyde by DERA (*E*_1_), and (iii) the conversion of acetaldehyde to Acetyl-CoA via AldDH (*E*_2_). Me-tabolite diffusion was described using a uniform diffusion coefficient, while metabolite transport across the cell membrane and AMC shell was defined by the boundary conditions. We assumed extracellular DNA enters the cell exclusively via active transport and cannot penetrate the AMC shell. The intermediate, Dr5P, was modeled to cross the cell membrane via both active transport and channel-mediated facilitated diffusion (concentration gradient-dependent passive transport through protein channels). In contrast, its transport across the AMC shell was modeled as permeability-dependent diffusion. Acetaldehyde was assumed to be passively transported across both the cell membrane and AMC shell.

For the AMC-present case, the variations in DNA 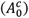, Dr5P 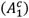 and acetaldehyde 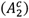 concentration in the cytosol are described as follows:

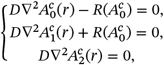

where *D* is the diffusion coefficient, *r* is the radial distance, and *R* is the reaction rate. *R* is determined using the Michaelis-Menten kinetic equation:

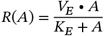

where *V*_*E*_ is the maximum reaction rate of the enzyme *E*, and *K*_*E*_ is the Michaelis constant.

The variations in Dr5P 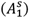 and acetaldehyde 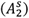 concentrations in the AMC are described as follows:

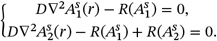

The boundary conditions at the cell membrane are:

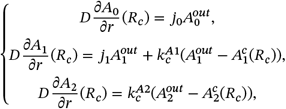

where *R*_*c*_ is cell radius, *j*_*i*_ is the active transport rate, 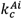 is the permea-bility of cell membrane to *A*_*i*_, and 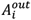is external concentration of *A*_*i*_.

The boundary conditions at the AMC shell interface are as follows:

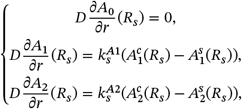

where *R*_*s*_ is AMC radius, 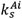 is the permeability of the AMC shell to *A*_*i*_.

The model for the AMC-absent case is formulated as follows:

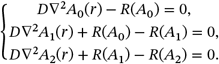

The boundary conditions at the cell membrane are:

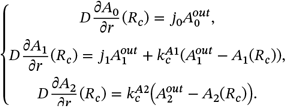

The resulting system of model equations was nondimensionalized and then solved numerically using a finite-difference scheme, implemented with the bvp5c solver in MATLAB. Further details regarding the solutions, simulation setup, and model parameter values are provided in Supplementary Note 1.

