## Supplementary Note 1 for "Microcompartments in archaeal ancestors of eukaryotes: a bioenergetic engine that could have fuelled eukaryogenesis"

### Supplementary Note 1: Mathematical modeling of metabolic fluxes for AMC-present and AMC-absent cases

#### 1 Model equations

This section presents the key governing equations for both the AMC-present and AMC-absent cases. For the AMC-present case, the variations in DNA ($A_0^c$), Dr5P ($A_1^c$) and acetaldehyde ($A_2^c$) concentrations in the cytosol are described as follows:

$$\begin{cases} D\nabla^2 A_0^c(r) - R(A_0^c) &= 0, & (1)\\ D\nabla^2 A_1^c(r) + R(A_0^c) &= 0, & (2)\\ D\nabla^2 A_2^c(r) &= 0, & (3)\end{cases}$$

where $D$ is the diffusion coefficient, $r$ is the radial distance, and $R$ is the reaction rate. $R$ is determined using the Michaelis-Menten kinetic equation:

$$R(A_0^c) = \frac{V_{E0}^c \cdot A_0^c}{K_{E0} + A_0^c}, \qquad (4)$$

where $V_{E0}^c$ is the maximum reaction rate of the enzyme $E_0$ (UPP), and $K_{E0}$ is the Michaelis constant. $V_{E0}^c$ can be calculated using the maximum reaction rate of enzyme active site $k_{E0}$ and the number of enzyme per cell $N_{E0}$ as follow:

$$V_{E0}^c = \frac{k_{E0} N_{E0}}{N_A \times (V_c - V_s)}, N_A = 6.022 \times 10^{23} mol^{-1}, \qquad (5)$$

where $V_c$ and $V_s$ respectively denote the cell and AMC volume.

The variations in Dr5P ($A_1^s$) and acetaldehyde ($A_2^s$) concentrations in the AMC are described as follows:

$$\begin{cases} D\nabla^2 A_1^s(r) - R(A_1^s) &= 0, & (6)\\ D\nabla^2 A_2^s(r) + R(A_1^s) - R(A_2^s) &= 0. & (7)\end{cases}$$

$R$ is determined as follow:

$$R(A_i^s) = \frac{V_{Ei}^s \cdot A_i^s}{K_{Ei} + A_i^s}, \quad i = 1, 2, \qquad (8)$$

where $E_1$ is DERA and $E_2$ is acetaldehyde, $V_{Ei}^s$ can be calculated as follow:

$$V_{Ei}^s = \frac{k_{Ei} N_{Ei}}{N_A \times V_s}, \quad i = 1, 2. \tag{9}$$

The boundary conditions at the cell membrane are as follows:

$$\begin{cases} D\dfrac{\partial A_0}{\partial r}(R_c) = j_0 A_0^{\text{out}}, & (10) \\ D\dfrac{\partial A_1}{\partial r}(R_c) = j_1 A_1^{\text{out}} + k_c^{A1}\left(A_1^{\text{out}} - A_1^c(R_c)\right), & (11) \\ D\dfrac{\partial A_2}{\partial r}(R_c) = k_c^{A2}\left(A_2^{out} - A_2^c(R_c)\right), & (12) \end{cases}$$

where $R_c$ is cell radius, $k_c^{Ai}$ is the permeability of cell membrane to $A_i$, and $A_i^{\text{out}}$ is external concentration of $A_i$.

The boundary conditions at the AMC shell interface are as follows:

$$\begin{cases} D\dfrac{\partial A_0}{\partial r}(R_s) = 0, & (13) \\ D\dfrac{\partial A_1}{\partial r}(R_s) = k_s^{A1}\left(A_1^c(R_s) - A_1^s(R_s)\right), & (14) \\ D\dfrac{\partial A_2}{\partial r}(R_s) = k_s^{A2}\left(A_2^c(R_s) - A_2^s(R_s)\right), & (15) \end{cases}$$

where $R_s$ is AMC radius, $k_s^{Ai}$ is the permeability of the AMC shell to $A_i$.

The model for the AMC-absent case is formulated as follows:

$$\begin{cases} D\nabla^2 A_0(r) - R(A_0) = 0, & (16) \\ D\nabla^2 A_1(r) + R(A_0) - R(A_1) = 0, & (17) \\ D\nabla^2 A_2(r) + R(A_1) - R(A_2) = 0, & (18) \end{cases}$$

$R$ is determined using the Michaelis-Menten kinetic equation:

$$R(A_i) = \frac{V_{Ei} \cdot A_i}{K_{Ei} + A_i}, \quad i = 0, 1, 2, \tag{19}$$

where

$$V_{Ei} = \frac{k_{Ei} N_{Ei}}{N_A \times V_c}, \quad i = 0, 1, 2. \tag{20}$$

The boundary conditions at the cell membrane are as follows:

$$\begin{cases} D\dfrac{\partial A_0}{\partial r}(R_c) = j_0 A_0^{\text{out}}, & (21) \\ D\dfrac{\partial A_1}{\partial r}(R_c) = j_1 A_1^{\text{out}} + k_c^{A1}\left(A_1^{\text{out}} - A_1(R_c)\right), & (22) \\ D\dfrac{\partial A_2}{\partial r}(R_c) = k_c^{A2}\left(A_2^{\text{out}} - A_2(R_c)\right). & (23) \end{cases}$$

#### 2 Solution for the AMC-present model

This section presents the numerical solution to the model describing metabolite concentrations in both the cytosol (governed by Eqs. (1) - (3)) and AMC (Eq. (6) and (7)), subject to the boundary conditions (Eq. (10) - (15)). First, we nondimensionalized the model with the following transformed variables:

$$
\begin{aligned}
&a_i = \frac{A_i}{K_{Ei}}, a_i^{\text{out}} = \frac{A_i^{\text{out}}}{K_{Ei}}, \rho = \frac{r}{R_c}, \rho_s = \frac{R_s}{R_c},\\
&\gamma_{12} = \frac{V_{E1}}{V_{E2}}, \kappa_{10} = \frac{K_{E1}}{K_{E0}}, D_{ai} = \frac{DK_{Ei}}{V_{Ei}R_c^2},\\
&B_{0c} = \frac{j_0 R_c}{D}, B_{1c} = \frac{j_1 R_c}{D}, B'_{1c} = \frac{k_c^{A1} R_c}{D},\\
&B_{2c} = \frac{k_c^{A2} R_c}{D}, B_{1s} = \frac{k_s^{A1} R_c}{D}, B_{2s} = \frac{k_s^{A2} R_c}{D}.
\end{aligned}
$$

The non-dimensional system includes the reaction-diffusion systems respectively in cytosol and in AMC:

$$
\begin{cases}
D_{a0}\dfrac{1}{\rho^2}\dfrac{\partial}{\partial\rho}(\rho^2\dfrac{\partial a_0^c}{\partial\rho}) = \dfrac{a_0^c}{1+a_0^c}, & (24)\\
D_{a0}\kappa_{10}\dfrac{1}{\rho^2}\dfrac{\partial}{\partial\rho}(\rho^2\dfrac{\partial a_1^c}{\partial\rho}) = -\dfrac{a_0^c}{1+a_0^c}, & (25)\\
\dfrac{1}{\rho^2}\dfrac{\partial}{\partial\rho}(\rho^2\dfrac{\partial a_2^c}{\partial\rho}) = 0, & (26)
\end{cases}
$$

$$
\begin{cases}
D_{a1}\dfrac{1}{\rho^2}\dfrac{\partial}{\partial\rho}(\rho^2\dfrac{\partial a_1^s}{\partial\rho}) = \dfrac{a_1^s}{1+a_1^s}, & (27)\\
D_{a2}\dfrac{1}{\rho^2}\dfrac{\partial}{\partial\rho}(\rho^2\dfrac{\partial a_2^s}{\partial\rho}) = \dfrac{a_2^s}{1+a_2^s} - \dfrac{a_1^s}{1+a_1^s}. & (28)
\end{cases}
$$

The boundary conditions at the cell membrane are transformed as follows:

$$
\begin{cases}
\dfrac{\partial a_0}{\partial\rho}(1) = B_{0c}a_0^{\text{out}}, & (29)\\
\dfrac{\partial a_1}{\partial\rho}(1) = B_{1c}a_1^{\text{out}} + B'_{1c}(a_1^{\text{out}} - a_1^c(1)), & (30)\\
\dfrac{\partial a_2}{\partial\rho}(1) = B_{2c}(a_2^{\text{out}} - a_2^c(1)). & (31)
\end{cases}
$$

The boundary conditions at the AMC shell interface are:

$$
\begin{cases}
\dfrac{\partial a_0}{\partial\rho}(\rho_s) = 0, & (32)\\
\dfrac{\partial a_1}{\partial\rho}(\rho_s) = B_{1s}(a_1^c(\rho_s) - a_1^s(\rho_s)), & (33)\\
\dfrac{\partial a_2}{\partial\rho}(\rho_s) = B_{2s}(a_2^c(\rho_s) - a_2^s(\rho_s)). & (34)
\end{cases}
$$

Due to the spherical symmetry, the boundary conditions at $r = 0$ (i.e., $\rho = 0$) should be:

$$\begin{cases} \dfrac{\partial a_1}{\partial \rho}(0) = 0, & (35) \\ \dfrac{\partial a_2}{\partial \rho}(0) = 0. & (36) \end{cases}$$

We then employed a finite-difference numerical scheme to solve the non-dimensionalized system of coupled second-order nonlinear ordinary differential equations in spherical coordinates. The boundary value problem (BVP) was resolved using **bvp5c** solver in MATLAB, a collocation-based routine that implements a finite-difference algorithm with adaptive mesh refinement. This approach provides accurate solutions for metabolite concentration profiles within the cytosol and AMC.

### 3 Solution for the AMC-absent model

The governing equations were transformed into dimensionless form using the following transformed variables:

$$a_i = \frac{A_i}{K_{Ei}}, \quad a_i^{\text{out}} = \frac{A_i^{\text{out}}}{K_{Ei}}, \quad \rho = \frac{r}{R_c},$$
$$\gamma_{01} = \frac{V_{E0}}{V_{E1}}, \quad \gamma_{12} = \frac{V_{E1}}{V_{E2}}, \quad D_{ai} = \frac{DK_{Ei}}{V_{Ei}R_c^2},$$
$$B_0 = \frac{j_0 R_c}{D}, \quad B_1 = \frac{j_1 R_c}{D}, \quad B_1' = \frac{k_c^{A1} R_c}{D}, \quad B_2 = \frac{k_c^{A2} R_c}{D}.$$

The non-dimensional equations are as follows:

$$\begin{cases} D_{a0}\dfrac{1}{\rho^2}\dfrac{\partial}{\partial \rho}(\rho^2 \dfrac{\partial a_0}{\partial \rho}) = \dfrac{a_0}{1 + a_0}, & (37) \\ D_{a1}\dfrac{1}{\rho^2}\dfrac{\partial}{\partial \rho}(\rho^2 \dfrac{\partial a_1}{\partial \rho}) = \dfrac{a_1}{1 + a_1} - \gamma_{01}\dfrac{a_0}{1 + a_0}, & (38) \\ D_{a2}\dfrac{1}{\rho^2}\dfrac{\partial}{\partial \rho}(\rho^2 \dfrac{\partial a_2}{\partial \rho}) = \dfrac{a_2}{1 + a_2} - \gamma_{12}\dfrac{a_1}{1 + a_1}. & (39) \end{cases}$$

with the boundary conditions respectively at cell membrane and at $\rho = 0$

$$\begin{cases} \dfrac{\partial a_0}{\partial \rho}(1) = B_0 a_0^{\text{out}}, & (40) \\ \dfrac{\partial a_1}{\partial \rho}(1) = B_1 a_1^{\text{out}} + B_1'(a_1^{\text{out}} - a_1(1)), & (41) \\ \dfrac{\partial a_2}{\partial \rho}(1) = B_2(a_2^{\text{out}} - a_2(1)), & (42) \end{cases}$$

$$\begin{cases} \dfrac{\partial a_0}{\partial \rho}(0) = 0, & (43) \\ \dfrac{\partial a_1}{\partial \rho}(0) = 0, & (44) \\ \dfrac{\partial a_2}{\partial \rho}(0) = 0. & (45) \end{cases}$$

The system was solved using **bvp5c** solver in MATLAB with a finite-difference numerical scheme as described above.

### 4 Simulations

The parameters were sourced from published literature, experimental measurements or were manually set in a reasonable range, as shown in Table S1. First, we performed simulations using the estimated parameters (as the baseline) and calculated the ratio of the flux through AldDH between the AMC and the cytosol. The flux through AldDH within AMC or cytosol was calculated using Eq. (46) and (47). The results indicated that the AldDH flux in the AMC-present case was approximatively 100-fold higher than in the AMC-free case.

$$f_{E2}^s = \frac{V_{E2}^s A_2^s}{K_{E2} + A_2^s} V_s. \tag{46}$$

$$f_{E2} = \frac{V_{E2} A_2}{K_{E2} + A_2} V_c. \tag{47}$$

To evaluate parameter sensitivity, we systematically increased and decreased each parameter by 10% and 20% from its baseline value for both the AMC-present (Fig. S1) and AMC-absent (Fig. S2) cases. The resulting fold changes relative to the baseline were documented. The AMC-present model exhibited higher sensitivity to variations in the extracellular DNA concentration ($A_0^{\text{out}}$), its active transport rate ($j_0$), the cell membrane permeability to Dr5P ($k_c^{A1}$), the AMC shell permeability to Dr5P ($k_s^{A1}$) and the AMC radius ($R_s$). In contrast, under AMC-absent conditions, in addition to $A_0^{\text{out}}$ and $j_0$, the kinetic parameters of AldDH ($k_{E2}$, $K_{E2}$ and $N_{E2}$) and the cell membrane permeability to acetaldehyde ($k_c^{A2}$) had a more substantial impact on the results. In both scenarios, the AldDH flux increased with higher substrate (DNA) uptake. The presence of the AMC led to a higher local concentration of AldDH and confined acetaldehyde more effectively within the compartment. As a result, the kinetic parameters of AldDH and the cell membrane permeability to acetaldehyde had a reduced influence in the AMC-present case. Conversely, since Dr5P is mainly produced in the cytosol, increased the AMC shell permeability elevated its concentration within the AMC, whereas higher cell membrane permeability reduced its cytosolic concentration and hence its concentration within AMC.

We further assessed the effect of parameter variations by applying large perturbations (upward and downward each value by up to 10-fold) in subsequent simulations, in order to confirm the enhancing effect of AMC on AldDH flux across a wide range of parameter values. The cell radius ($R_c \in [0.3\mu m, 0.9\mu m]$) and AMC radius ($R_s[0.025\mu m, 0.1\mu m]$) were excluded from the ten-fold adjustment and were instead varied within their experimentally measured ranges. The results were in the main text.

Table S1. Model parameter description and estimated value

| Parameter | Description | Estimated value | Unit | Source |
|---|---|---|---|---|
| $D$ | Diffusivity of metabolites in the cellular milieu | $1.19 \times 10^6$ | $\mu m^2/h$ | [1] |
| $k_{E_0}$ | Maximum reaction rate of UPP | $1.365 \times 10^5$ | $1/h$ | [2] |
| $K_{E_0}$ | Michaelis-Menten constant of UPP | 170 | $\mu M$ | [2] |
| $N_{E_0}$ | Number of UPP per cell | 1000 | $1/cell$ | Manually set |
| $k_{E_1}$ | Maximum reaction rate of DERA | $1.44 \times 10^5$ | $1/h$ | [3] |
| $K_{E_1}$ | Michaelis-Menten constant of DERA | 290 | $\mu M$ | [3] |
| $N_{E_1}$ | Number of DERA per cell | 1000 | $1/cell$ | Manually set |
| $k_{E_2}$ | Maximum reaction rate of AldDH | $3.9 \times 10^4$ | $1/h$ | [4] |
| $K_{E_2}$ | Michaelis-Menten constant of AldDH | 1000 | $\mu M$ | [4] |
| $N_{E_2}$ | Number of AldDH per cell | 2500 | $1/cell$ | [5] |
| $j_0$ | Active transport rate of DNA | 100 | $\mu m/h$ | Manually set |
| $j_1$ | Active transport rate of Dr5P | 100 | $\mu m/h$ | Manually set |
| $k_c^{A1}$ | Cell membrane permeability to Dr5P by channel-mediated facilitated diffusion | 36 | $\mu m/h$ | Manually set |
| $k_c^{A2}$ | Cell membrane permeability to acetaldehyde | $3.6 \times 10^4$ | $\mu m/h$ | [6] |
| $k_s^{A1}$ | AMC shell permeability to Dr5P | $3.6 \times 10^2$ | $\mu m/h$ | [7] |
| $k_s^{A2}$ | AMC shell permeability to acetaldehyde | $3.6 \times 10^2$ | $\mu m/h$ | [7] |
| $A_0^{out}$ | External concentration of DNA | 10 | $\mu M$ | Manually set |
| $A_1^{out}$ | External concentration of Dr5P | 0.01 | $\mu M$ | Manually set |
| $A_2^{out}$ | External concentration of acetaldehyde | 0 | $\mu M$ | Manually set |
| $R_s$ | AMC radius | 0.04 | $\mu m$ | Measured |
| $R_c$ | Cell radius | 0.6 | $\mu m$ | [8] |

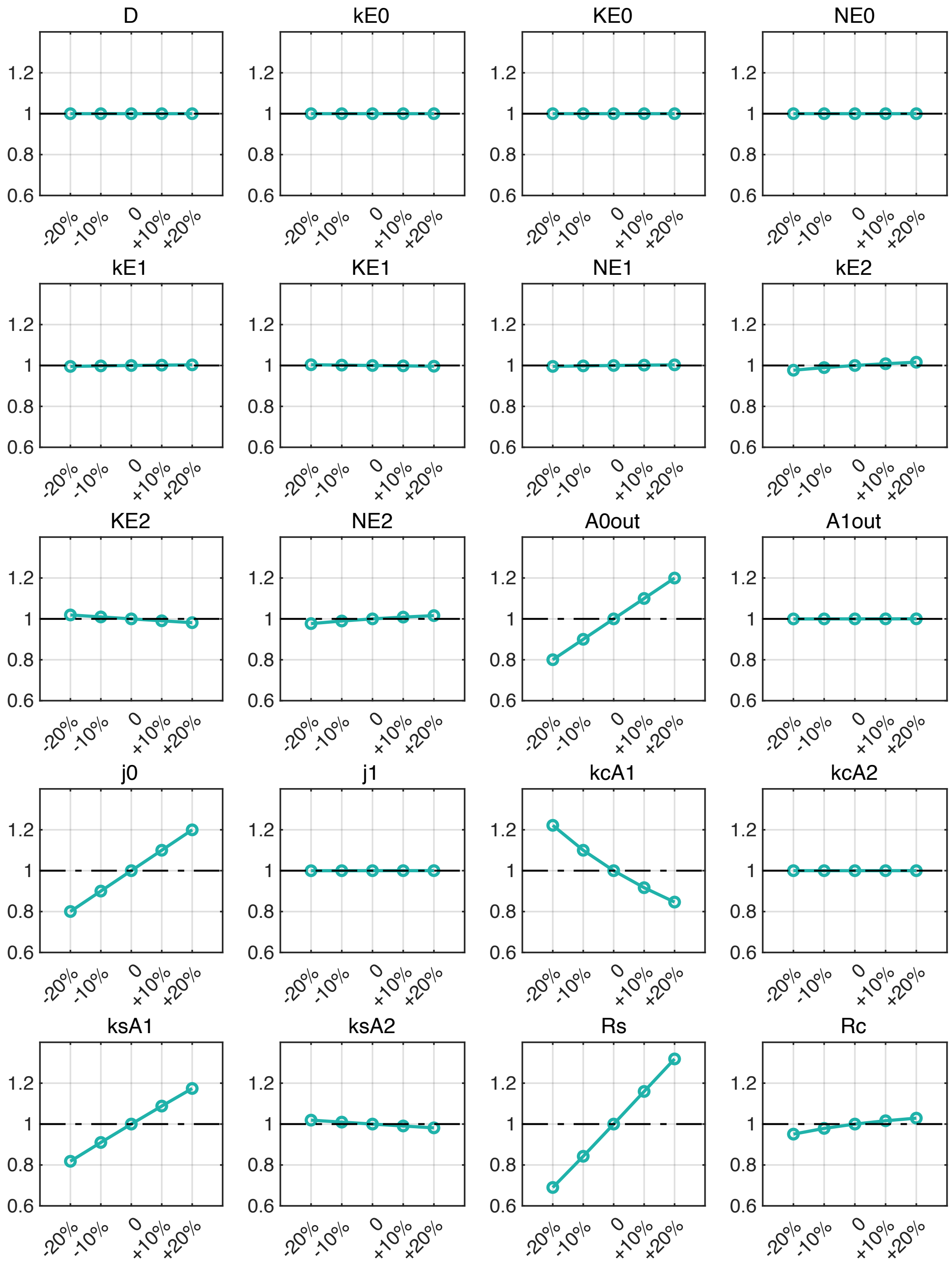


Fig. S1. **Sensitivity of AldDH flux to model parameter variation in the presence of AMC.** The x-axis represents the percentage change (increase or decrease) in each parameter relative to its baseline value; the y-axis represents the ratio of the perturbed AldDH flux to the baseline flux.

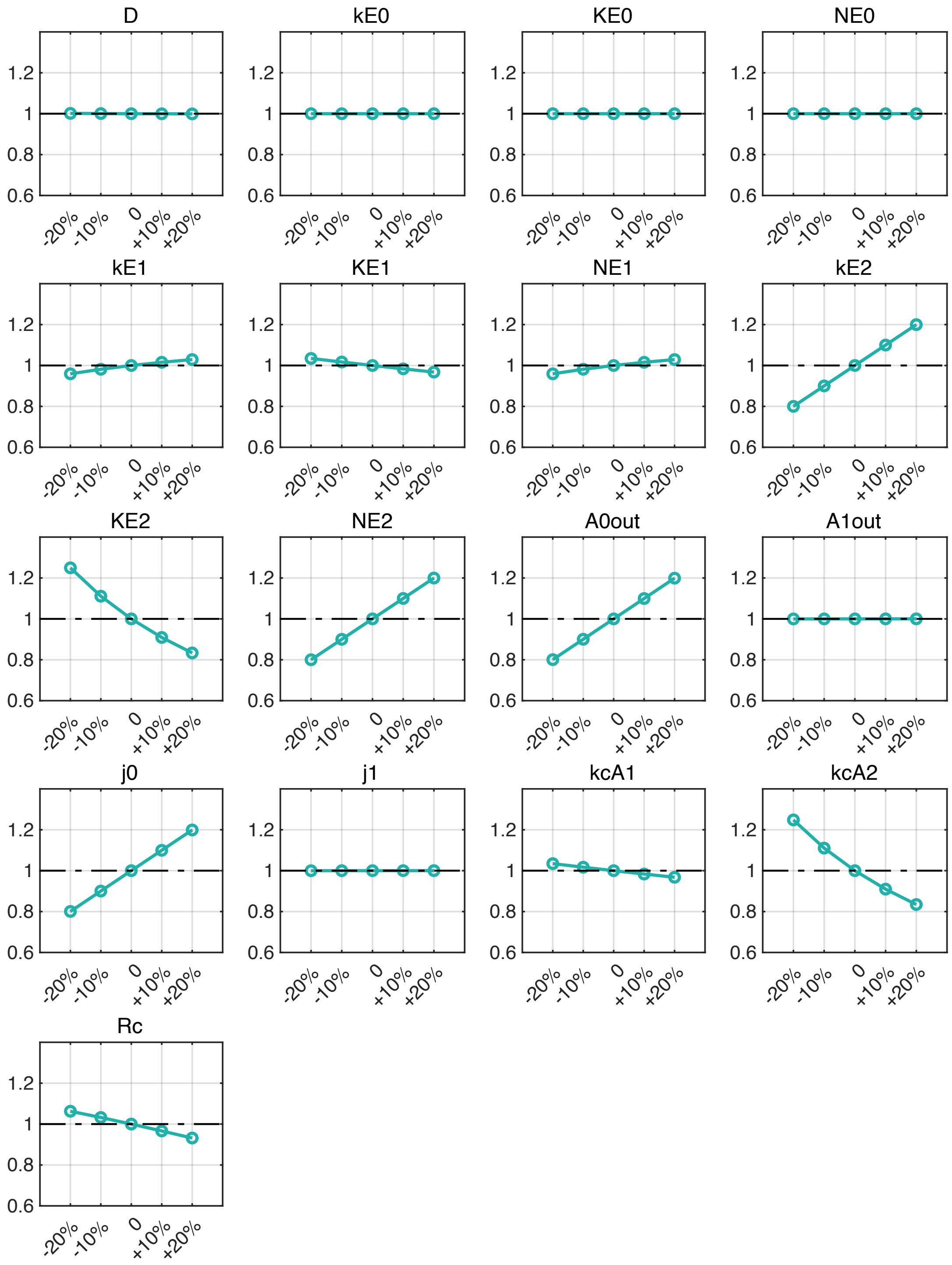


Fig. S2. **Sensitivity of AldDH flux to model parameter variation in the absence of AMC.** The x-axis represents the percentage change (increase or decrease) in each parameter relative to its baseline value; the y-axis represents the ratio of the perturbed AldDH flux to the baseline flux.
