## Supplementary Figures for "Microcompartments in archaeal ancestors of eukaryotes: a bioenergetic engine that could have fuelled eukaryogenesis"

bioRχiv

**Huan Du,[a,#] Aoxiang Xu,[a,#] Xiaoyuan Feng,[a,#] Wen-cong Huang,[a,e] Huayu Li,[a] Li Liu,[a] Yuling Li,[a] Siyu Zhang,[a] Ning Song,[a] Kathryn E. Appler,[b] Brett J. Baker,[b,c] Eugene V. Koonin,[d] Meng Li,[a,*] Yang Liu,[a,*]**
[a]Archaeal Biology Center, Synthetic Biology Research Center, Shenzhen Key Laboratory of Marine Microbiome Engineering, Key Laboratory of Marine Microbiome Engineering of Guangdong Higher Education Institutes, Institute for Advanced Study, Shenzhen University, Shenzhen 518060, China
[b]Department of Marine Science, University of Texas at Austin, Marine Science Institute, Port Aransas, TX, USA
[c]Department of Integrative Biology, University of Texas at Austin, Austin, TX, USA
[d]Computational Biology Branch, Division of Intramural Research, National Library of Medicine, Bethesda, MD 20894, USA
[e]Present address: Royal Netherlands Institute for Sea Research, Netherlands

[#]These authors contributed equally to this work

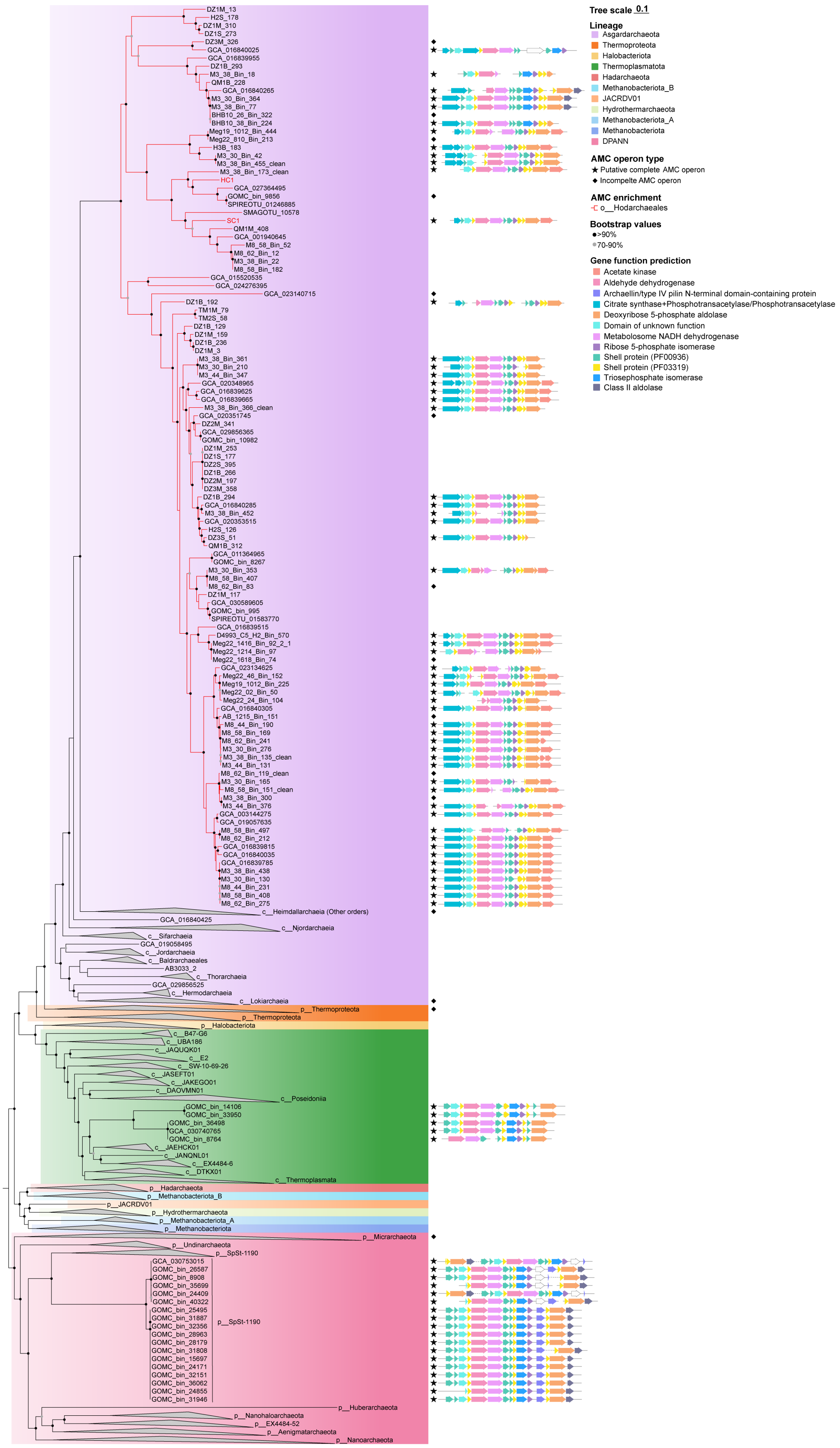


**Supplementary Fig. 1 | Phylogenetic distribution of the AMC operon in Archaea, with a detailed view of the Hodarchaeales lineages.** A maximum-likelihood phylogenetic tree of 53 archaeal marker gene concatenation was reconstructed from 1470 representative archaeal genomes using IQ-Tree v2.3.6 (-m LG+C60+G -mwopt -B 1000 -alrt 1000) with the posterior mean site frequency approximation using the guide tree was inferred under LG+G model. Branches corresponding to the order Hodarchaeales are marked in red. The two isolated Hodarchaeales strains, SC1 and HC1, are specifically labeled in red. Putative complete AMC operons are annotated alongside their corresponding genomes (right panel).

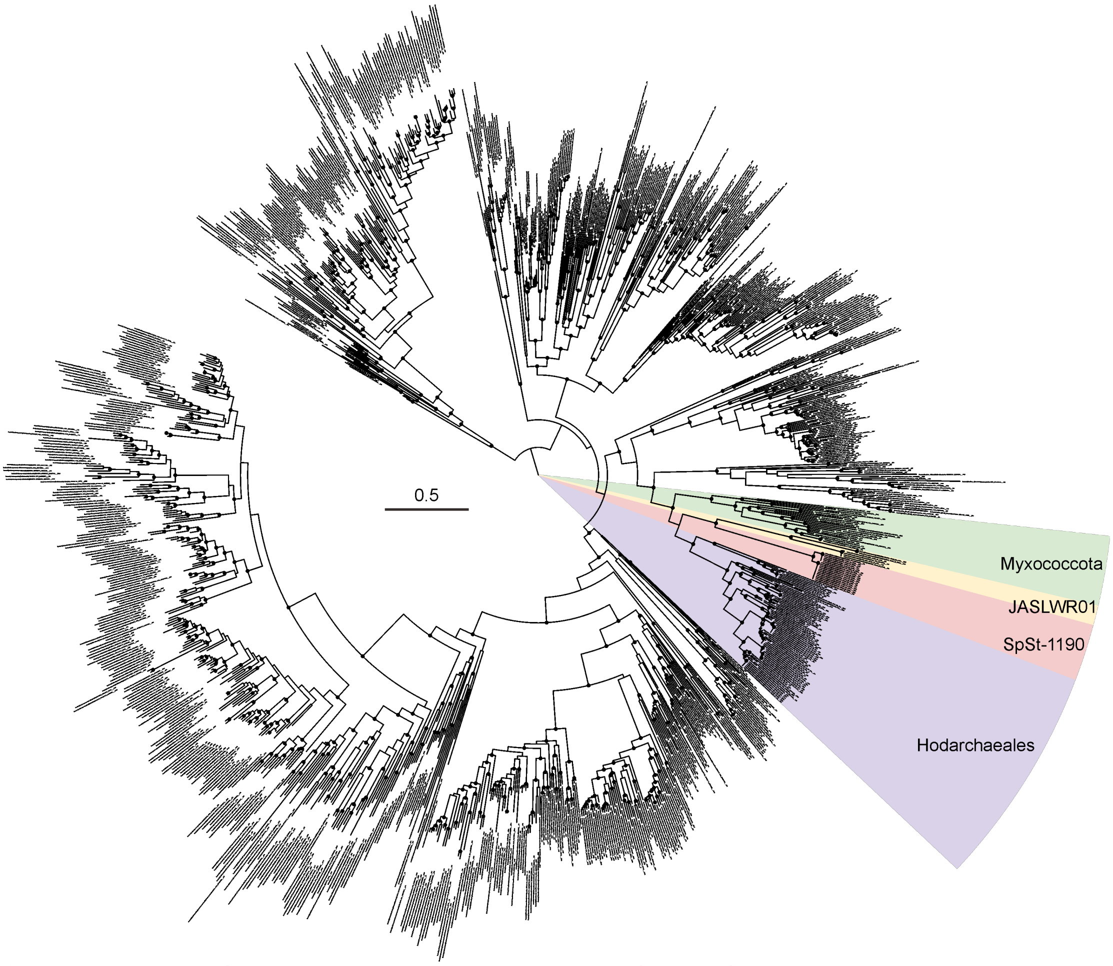


**Supplementary Fig. 2 | Phylogenetic analysis of ALDH protein (PF00171).**
Maximum likelihood phylogenetic tree of aldehyde dehydrogenase (ALDH) proteins based on the Pfam family PF00171. The scale bar representing 0.5 substitutions per site. Black circles at nodes indicate bootstrap support values larger than 0.8.

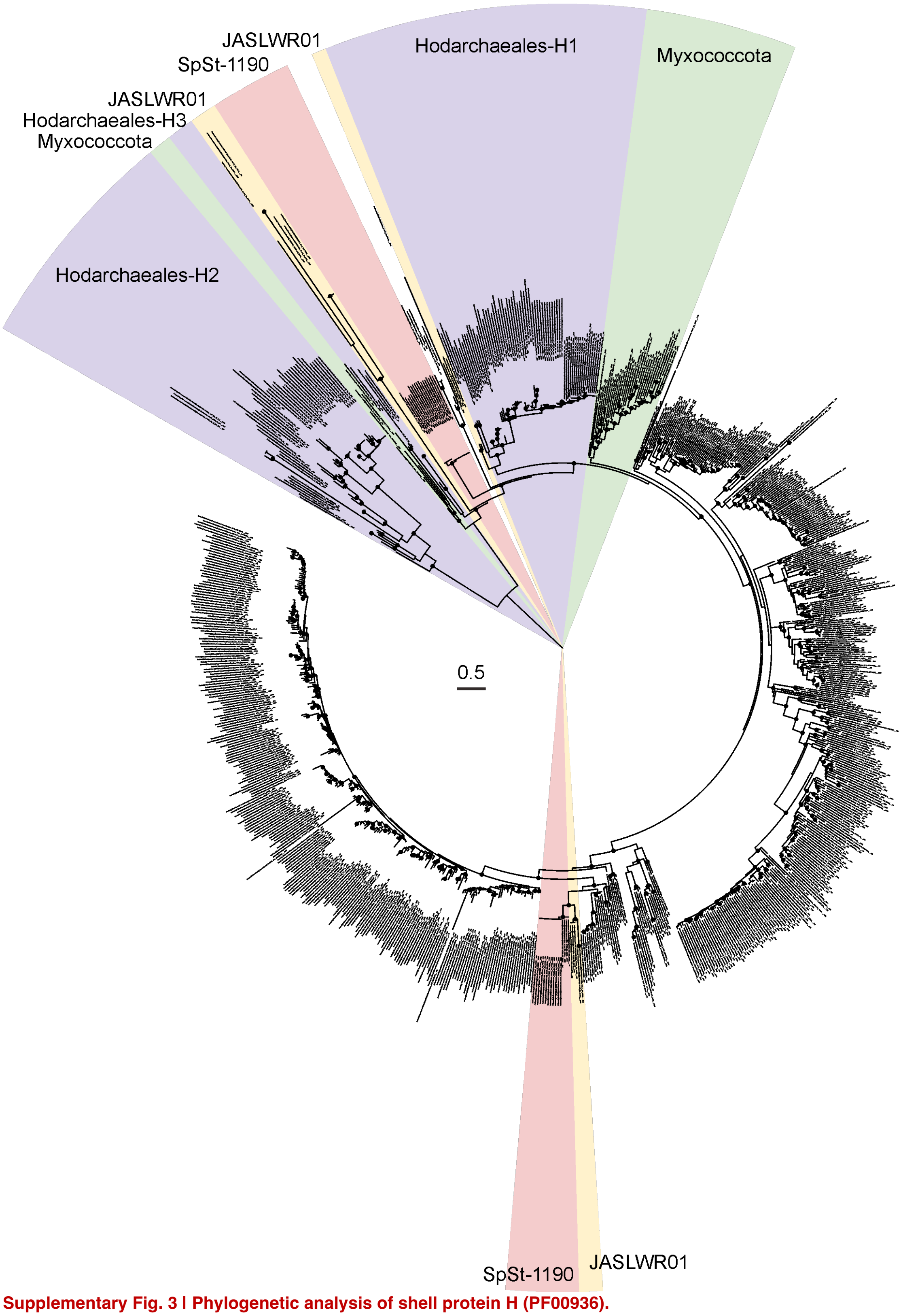


**Supplementary Fig. 3 | Phylogenetic analysis of shell protein H (PF00936).**
Maximum likelihood phylogenetic tree of shell protein H based on the Pfam family PF00936 and https://www.kerfeldlab.org/bmc-locus-hmms.html. The scale bar representing 0.5 substitutions per site. Black circles at nodes indicate bootstrap support values larger than 0.8.

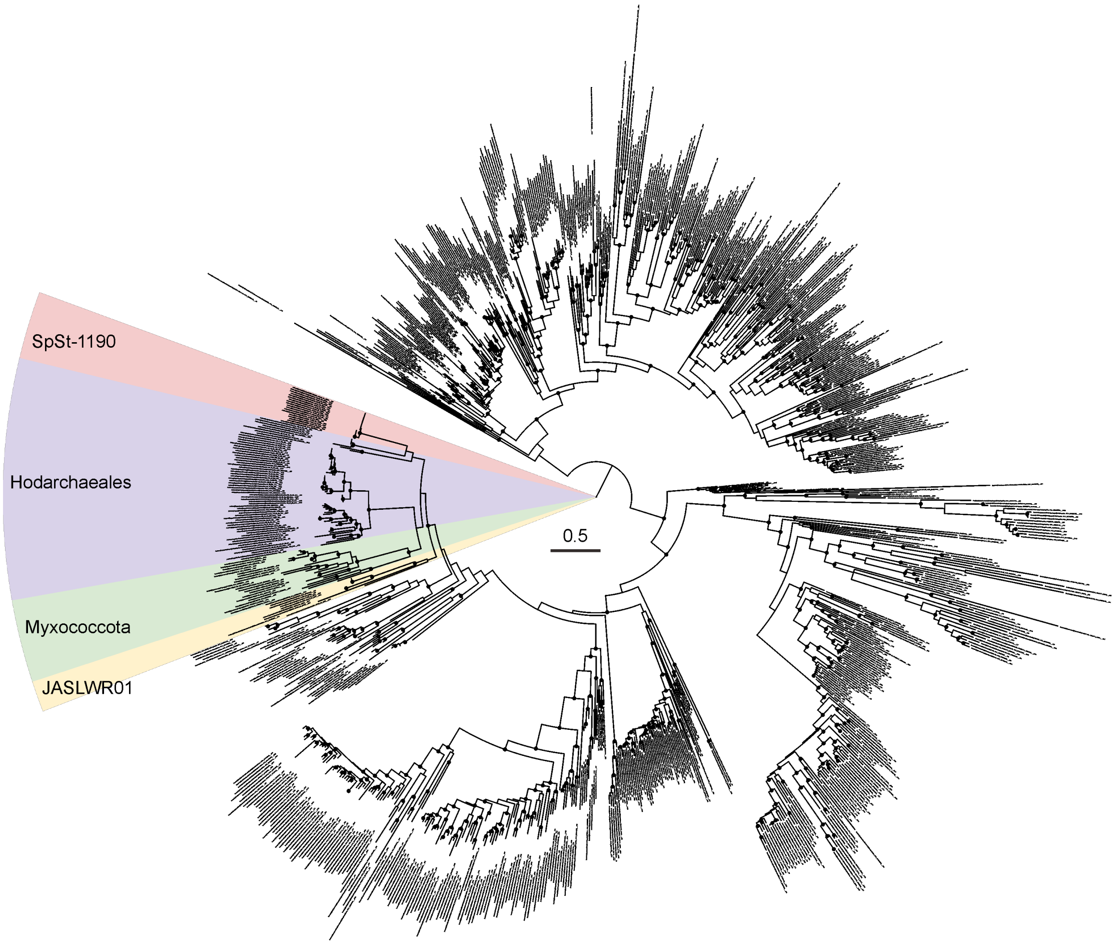


**Supplementary Fig. 4 | Phylogenetic analysis of shell protein Ts (PF00936).**
Maximum likelihood phylogenetic tree of BMC domain proteins based on the Pfam family PF00936 and https://www.kerfeldlab.org/bmc-locus-hmms.html. The scale bar representing 0.5 substitutions per site. Black circles at nodes indicate bootstrap support values larger than 0.8.

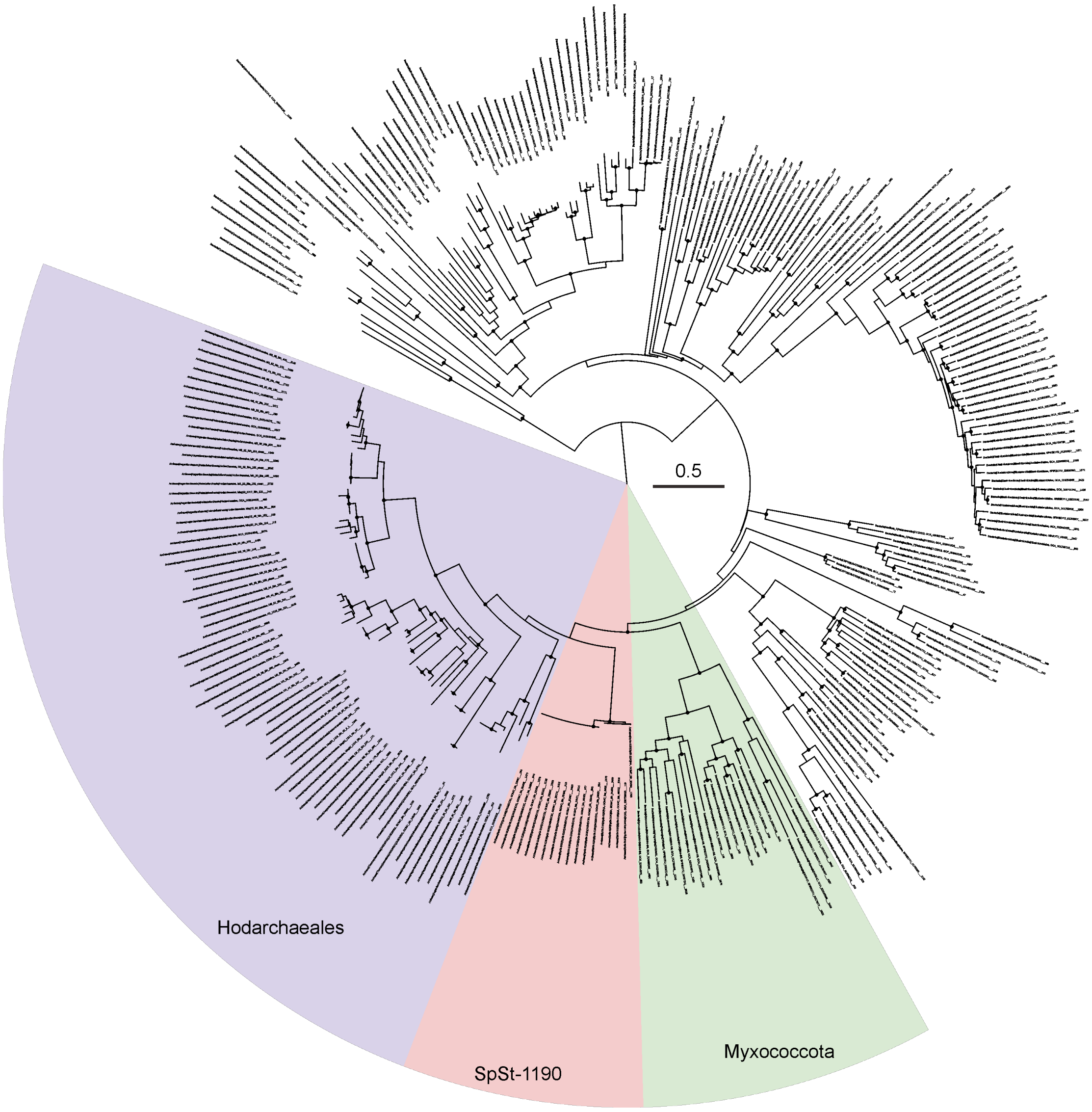


**Supplementary Fig. 5 | Phylogenetic analysis of Metabolosome Nicotinamide Adenine Dinucleotide Hydrogen (NADH) dehydrogenase (PF01512).**
Maximum likelihood phylogenetic tree of MNdh based on the Pfam family PF01512. The scale bar representing 0.5 substitutions per site. Black circles at nodes indicate bootstrap support values larger than 0.8.

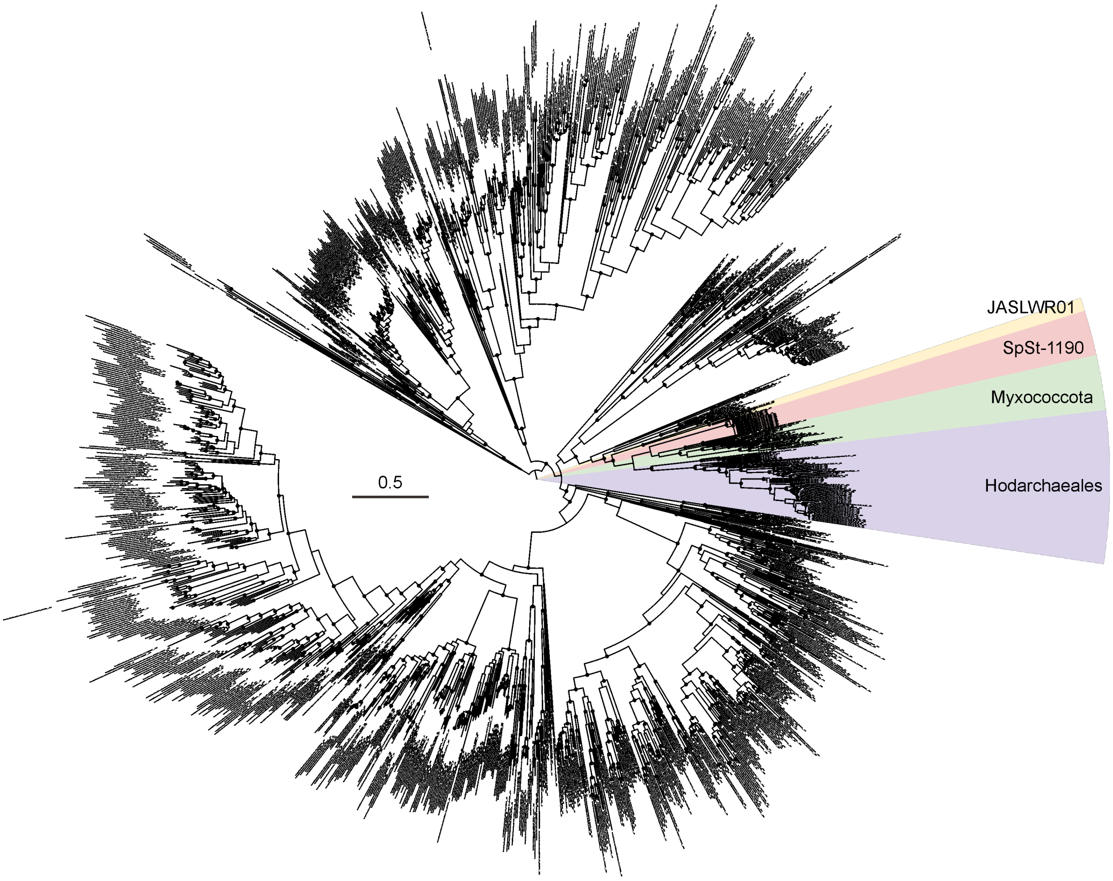


**Supplementary Fig. 6 | Phylogenetic analysis of deoxyribose 5-phosphate aldolase (PF01791).**
Maximum likelihood phylogenetic tree of DERA based on the Pfam family PF01791. The scale bar representing 0.5 substitutions per site. Black circles at nodes indicate bootstrap support values larger than 0.8.

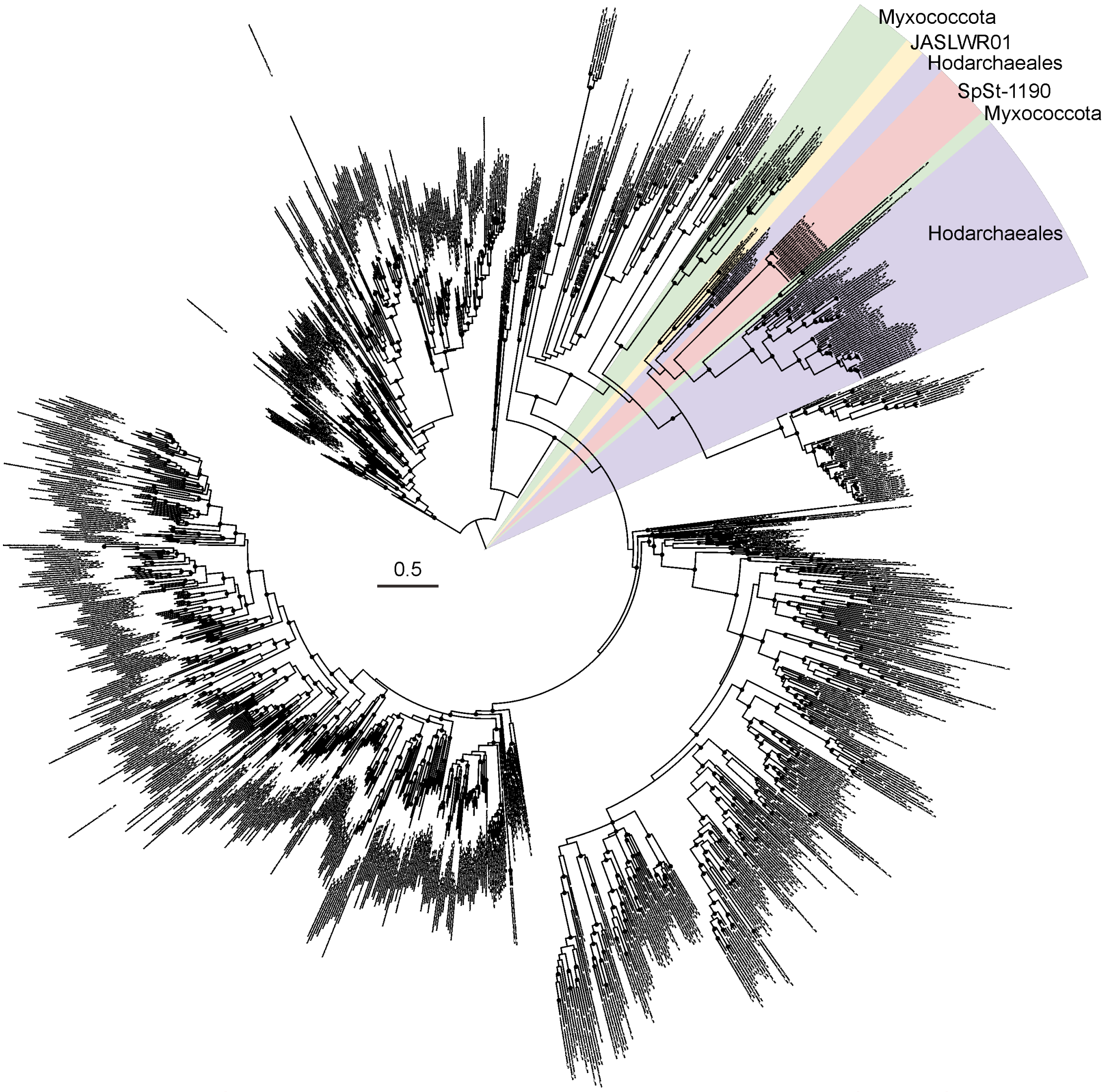


**Supplementary Fig. 7 | Phylogenetic analysis of Ribose sugar-phosphate Isomerase (PF02502).**
Maximum likelihood phylogenetic tree of RPI based on the Pfam family PF02502. The scale bar representing 0.5 substitutions per site. Black circles at nodes indicate bootstrap support values larger than 0.8.

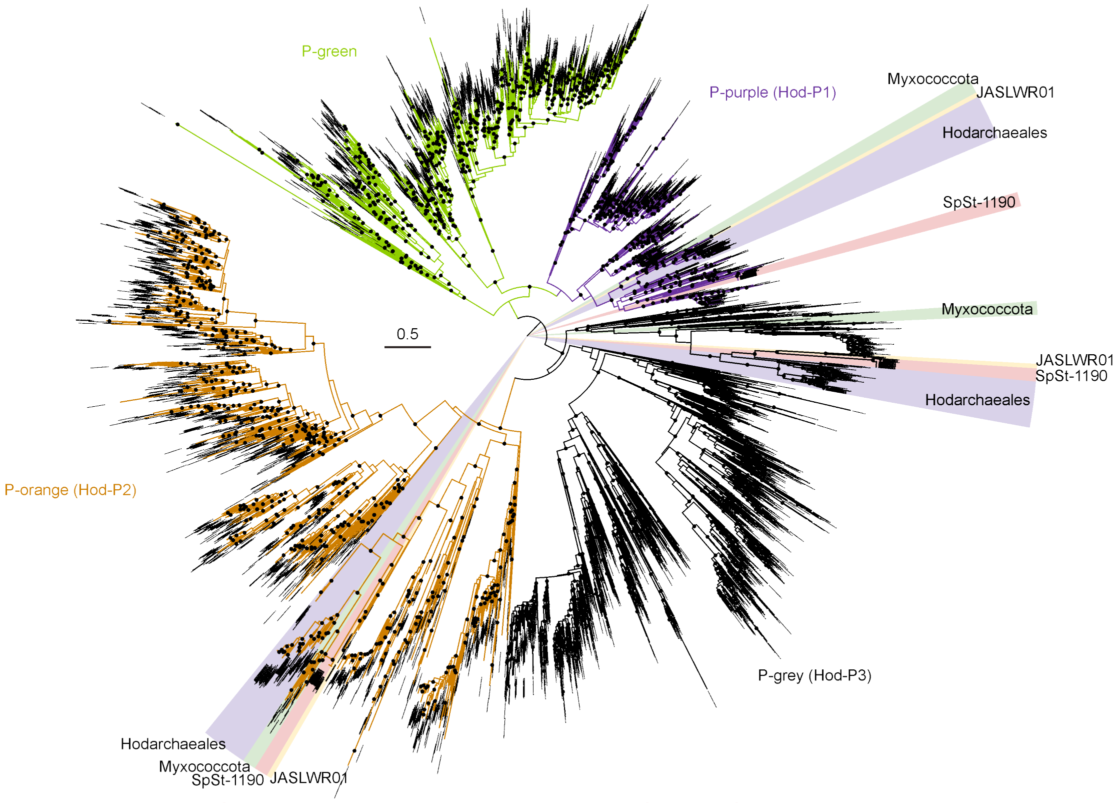


**Supplementary Fig.8 l Phylogenetic analysis of shell protein P (PF03319).**
Maximum likelihood phylogenetic tree of shell protein P based on the Pfam family PF03319 and https://www.kerfeldlab.org/bmc-locus-hmms.html. The scale bar representing 0.5 substitutions per site. Black circles at nodes indicate bootstrap support values larger than 0.8.

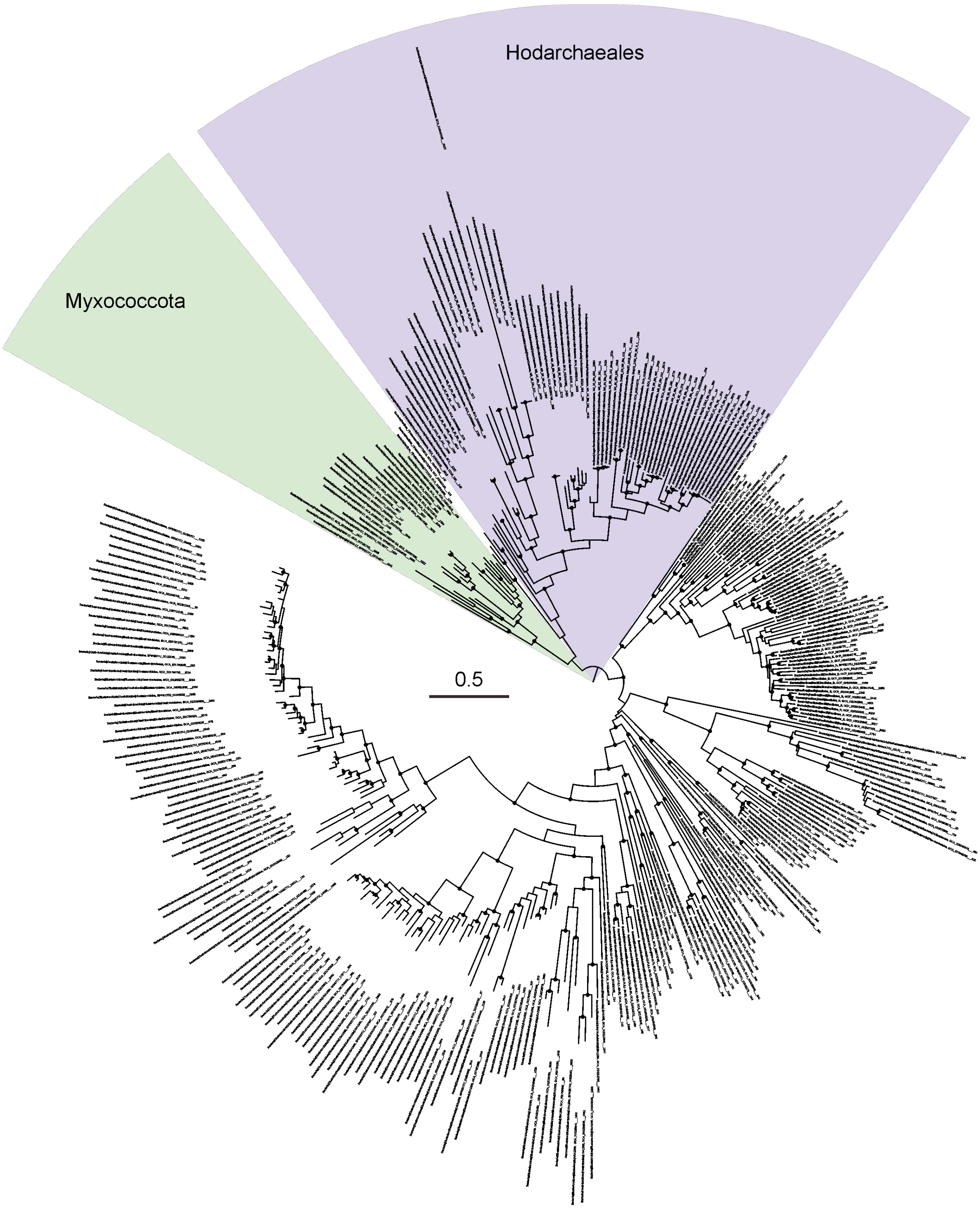


**Supplementary Fig. 9 | Phylogenetic analysis of phosphotransacylase (PF06130).**
Maximum likelihood phylogenetic tree of PTAC based on the Pfam family PF06130. The scale bar representing 0.5 substitutions per site. Black circles at nodes indicate bootstrap support values larger than 0.8.

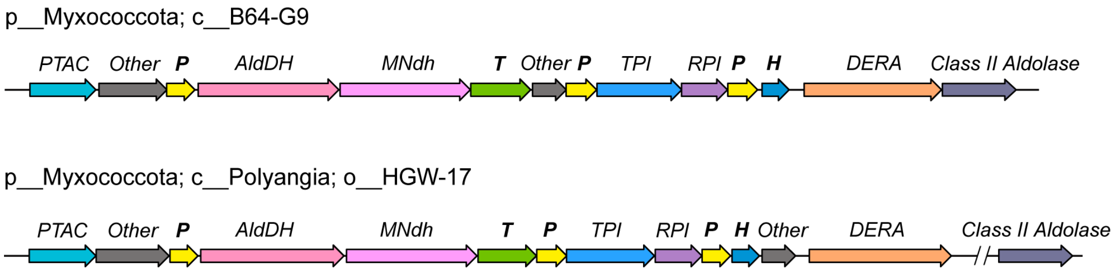


**Supplementary Fig. 10 | Gene organization of BMC operons in Myxococcota lineages (class B64-G9 and order HGW-17).**
The BMC operons are from the representative genomes in the class B64-G9 (GCA_022710975) and the order HGW-17 (GCA_016937155) in Myxococcota. The operons consisting of genes encoding five structural components (three BMC-Ps, one BMC-H and one BMC-T) marked in bold along with genes encoding metabolic enzymes. Abbreviations include PTAC (phosphotransacetylase), AldDH (acetaldehyde dehydrogenase), NADH dehydrogenase (MNdh), TPI (triosephosphate isomerase), RPI (ribose 5-phosphate isomerase), DERA (deoxyribose 5-phosphate aldolase).

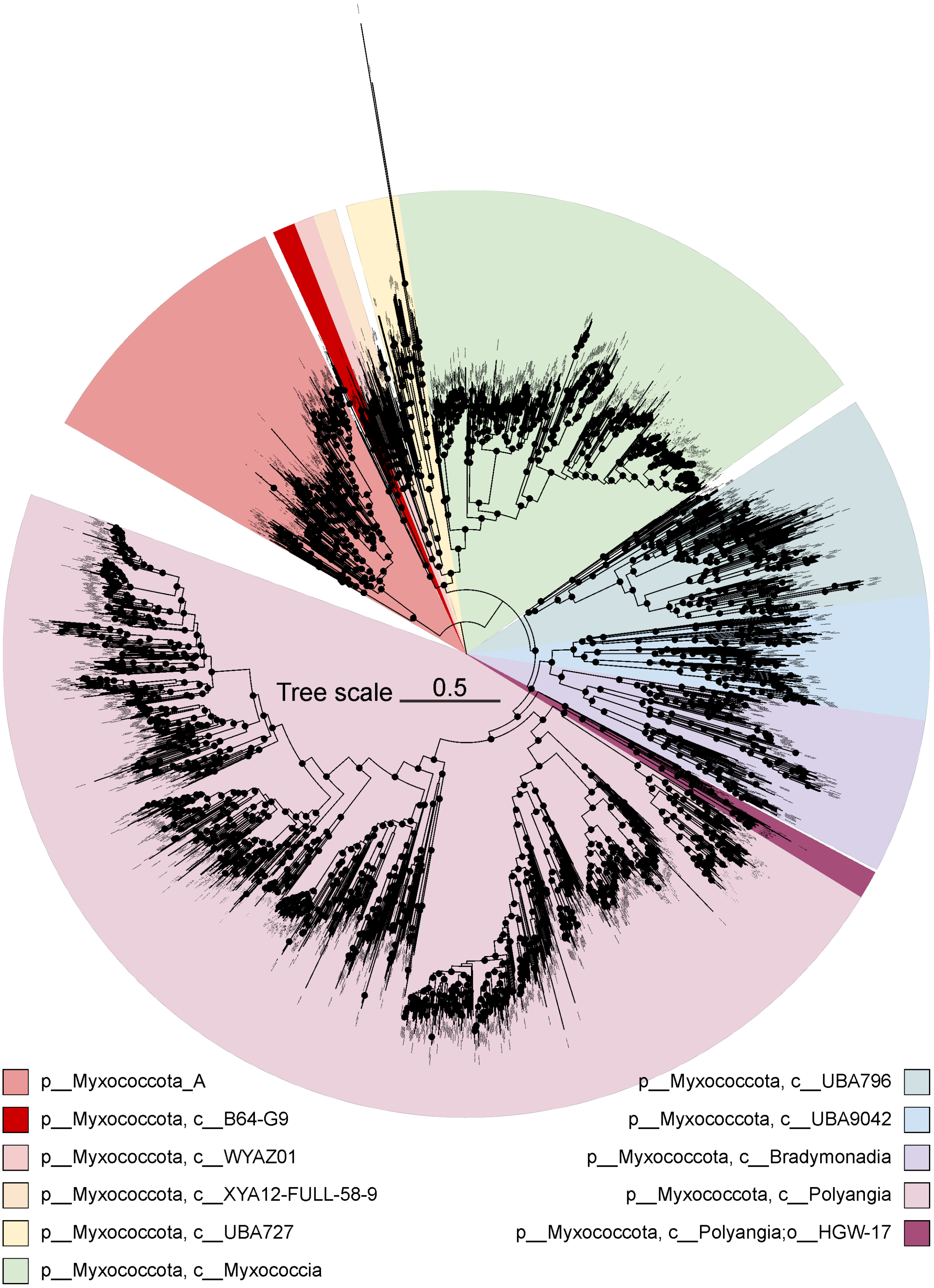


**Supplementary Fig. 11 | Species tree of Myxococcota showing the position of B64-G9 class and HGW-17 order.**
The phylogenetic tree was constructed using IQ-tree with LG+I+G model based on 120 conserved bacterial proteins following the GTDB-tk pipeline. Topology of genomes from GTDB was used as a guide tree. Nodes with SH-aLRT bootstrap values greater than 95% are indicated by solid circles.

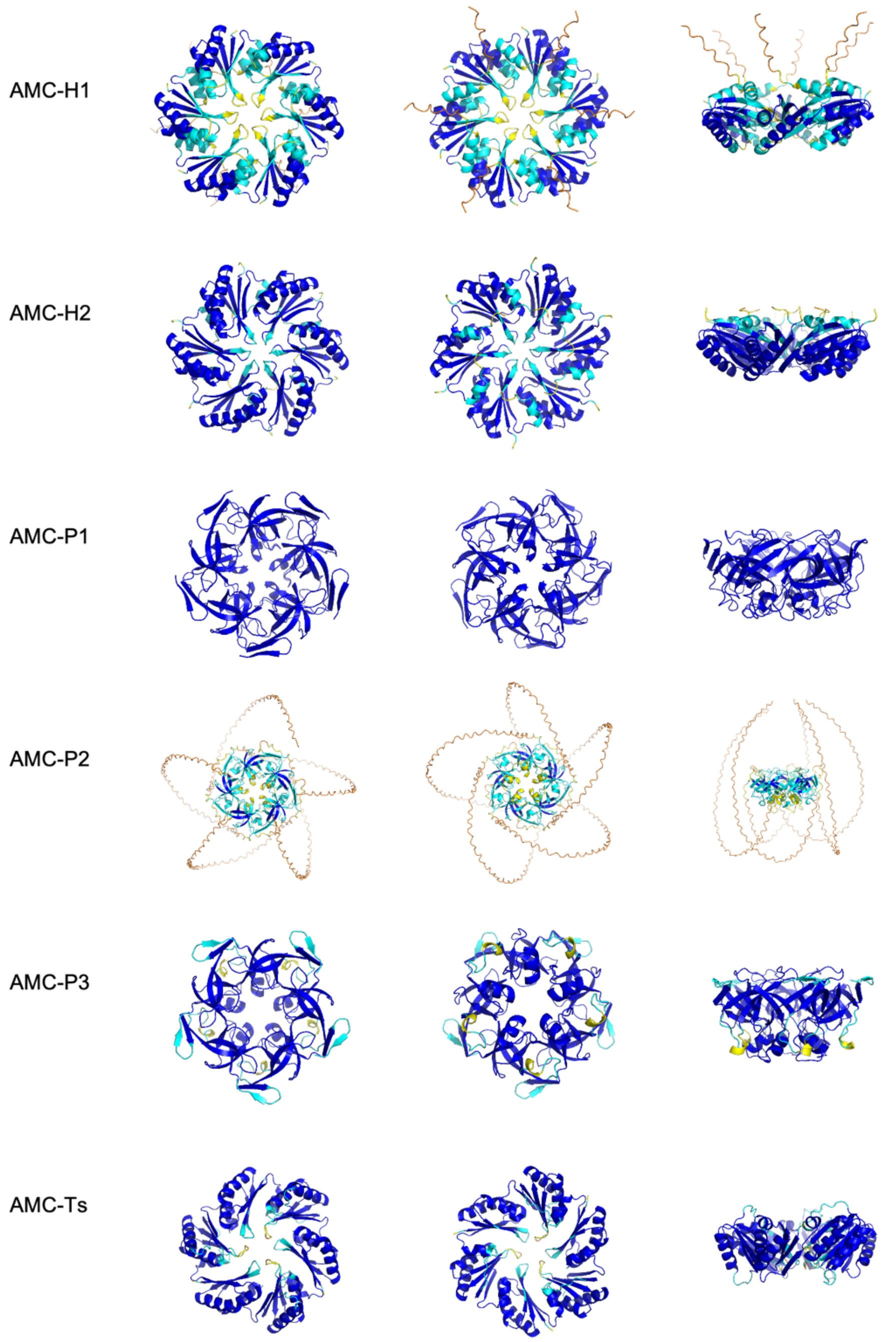


**Supplementary Fig. 12 | Structural models of shell proteins.**
AlphaFold3 model of six types of AMC shell protein structures from Hodarchaeales (accession number: GCA_003144275) with top, bottom and side views. Colors represent the secondary structure confidence (pLDDT) (Blue = very high, cyan = confident, yellow = low, orange = very low).

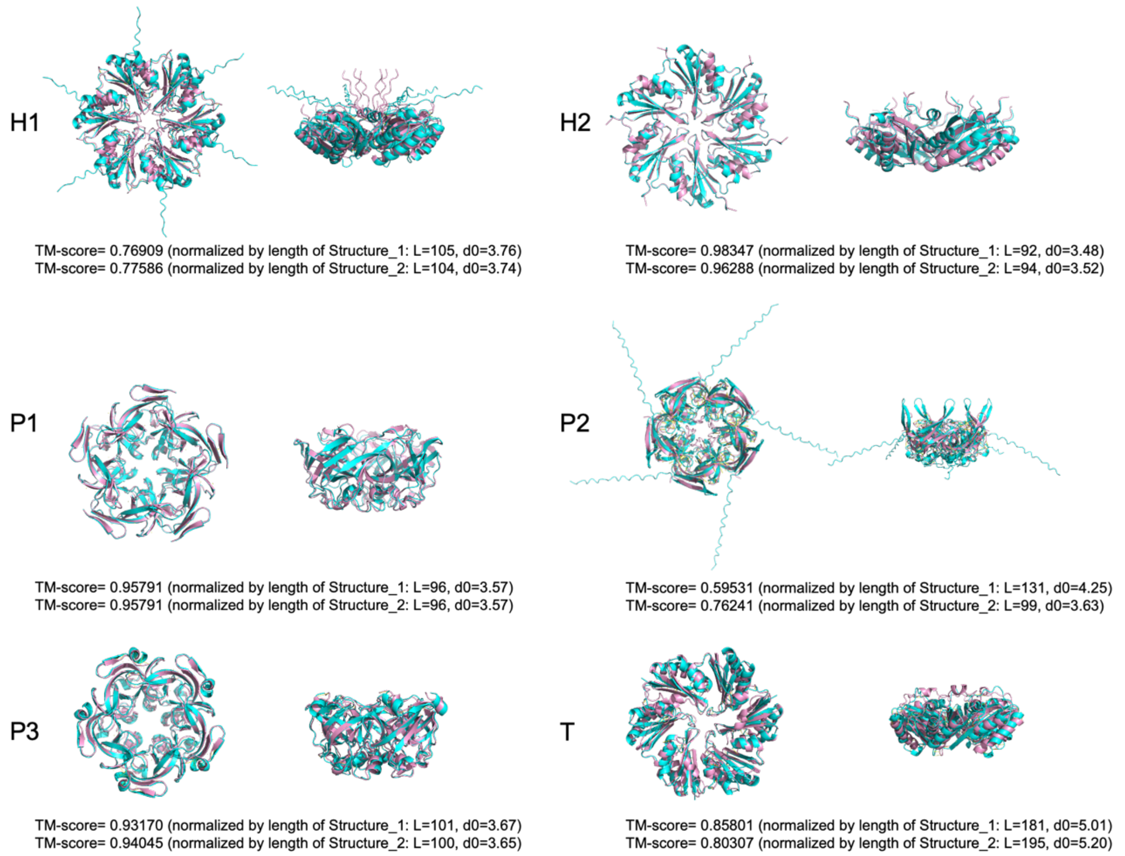


**Supplementary Fig. 13 | Alignment of the AMC shell protein structures from Hodarchaeales with those of BMC from Myxococcota.** Shown in cyan is the Alphafold3 model of the AMC shell proteins (Structure 1) from Hodarchaeales strain SC1; in pink is the Alphafold3 model of the BMC shell proteins (Structure 2) from Myxococcota B64-G9 (accession number: GCA_003647095). The TM-score was computed using US-align[5].

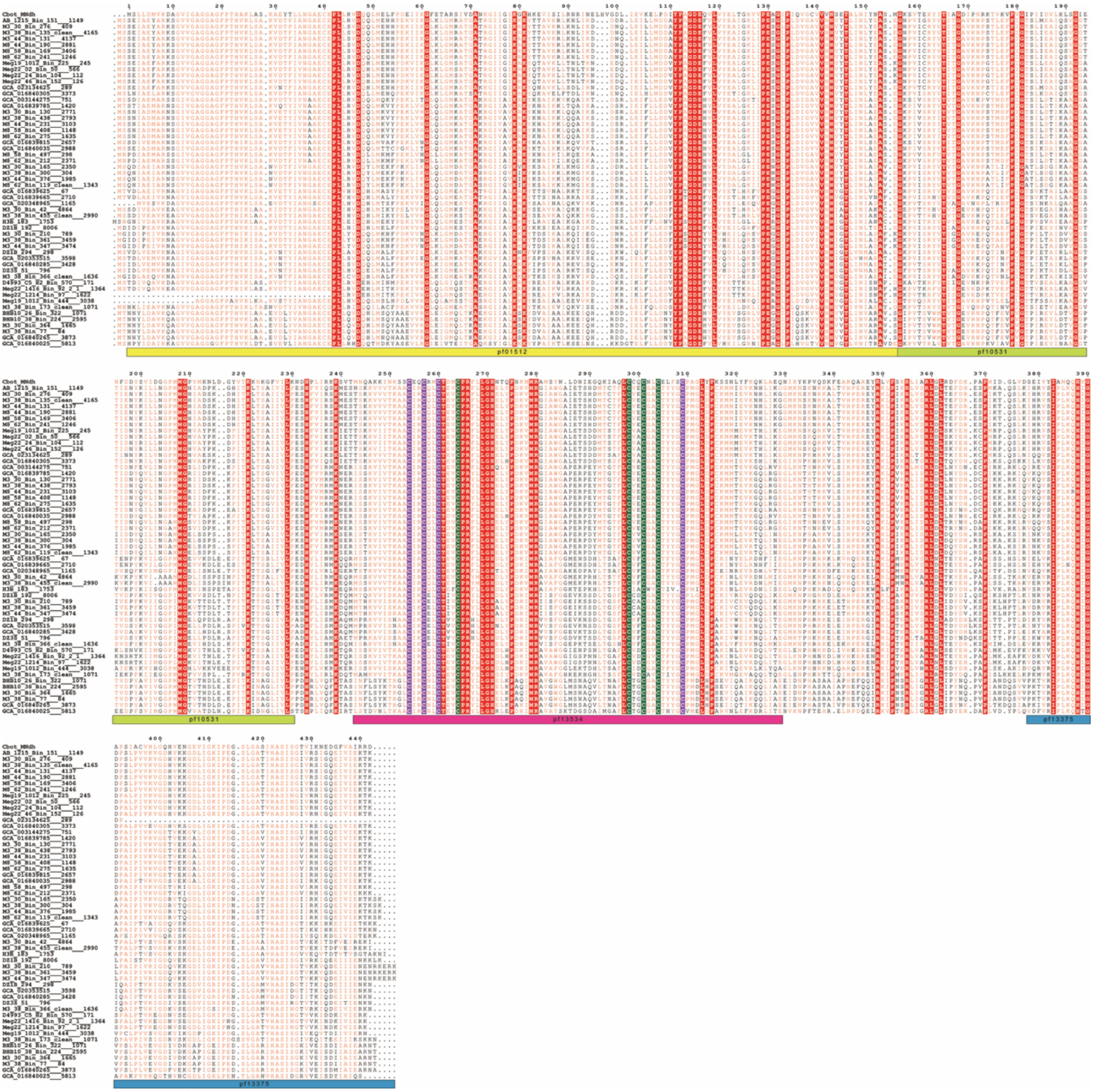

**Supplementary Fig. 14 | Sequence alignment of AMC MNdh with MNdh from the GRM1 operon of *Clostridium botulinum*[1].**
Cysteines coordinating the Fe-S clusters are colored in green and purple for the two different 4Fe-4S clusters, respectively, while completely conserved residues are colored in red. Multiple sequence alignment was performed using MAFFT 7.0[2] and visualized with ESPript 3.0[3].

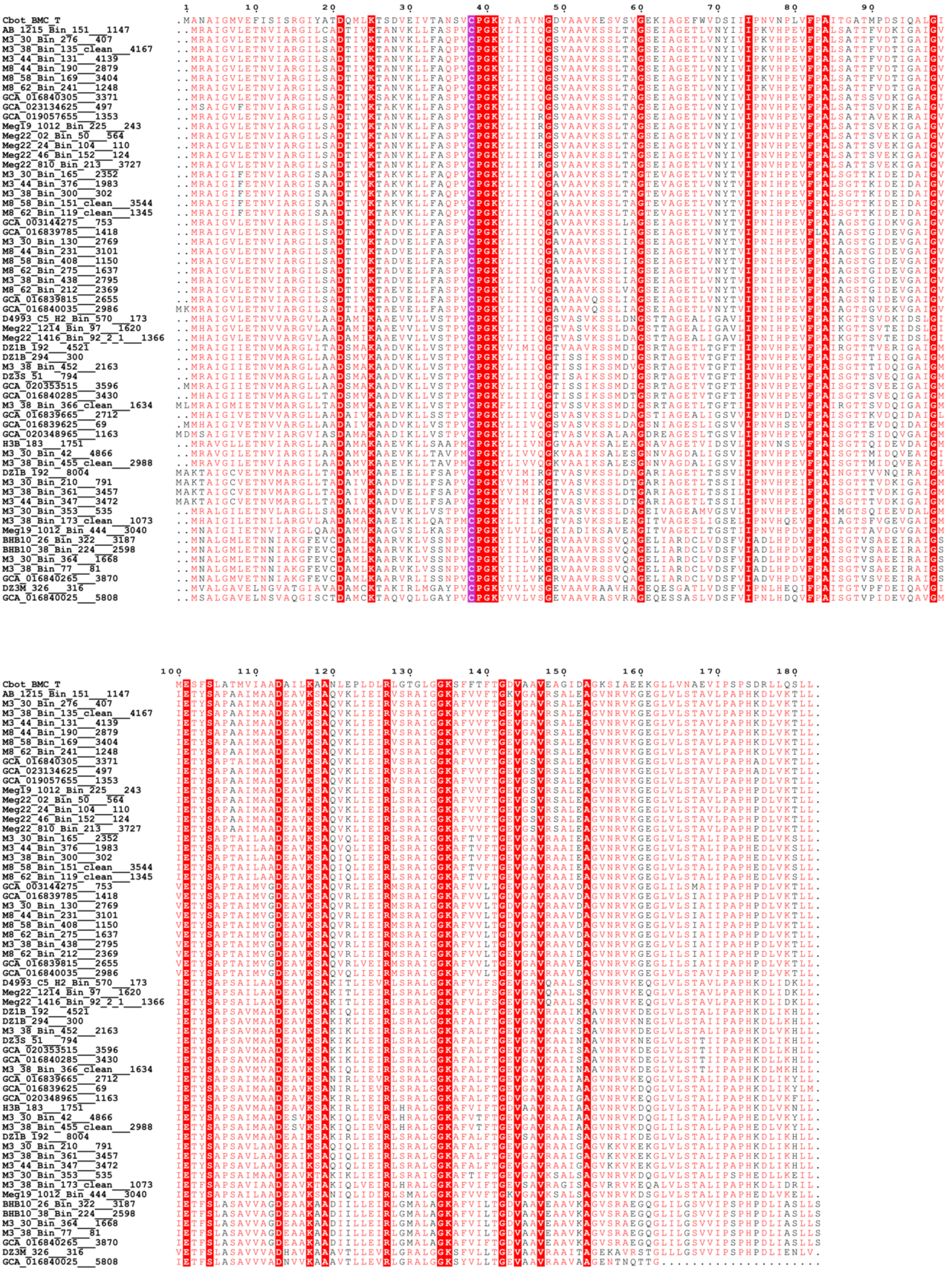

**Supplementary Fig. 15 | Sequence alignment of AMC-T$^{se}$ with BMC-T$^{se}$ from the GRM1 operon of Clostridium botulinum.** Cysteine coordinating the Fe-S cluster is colored in purple, while completely conserved residues are colored in red. Multiple sequence alignment was performed using MAFFT 7.0[2] and visualized with ESPript 3.0[3].

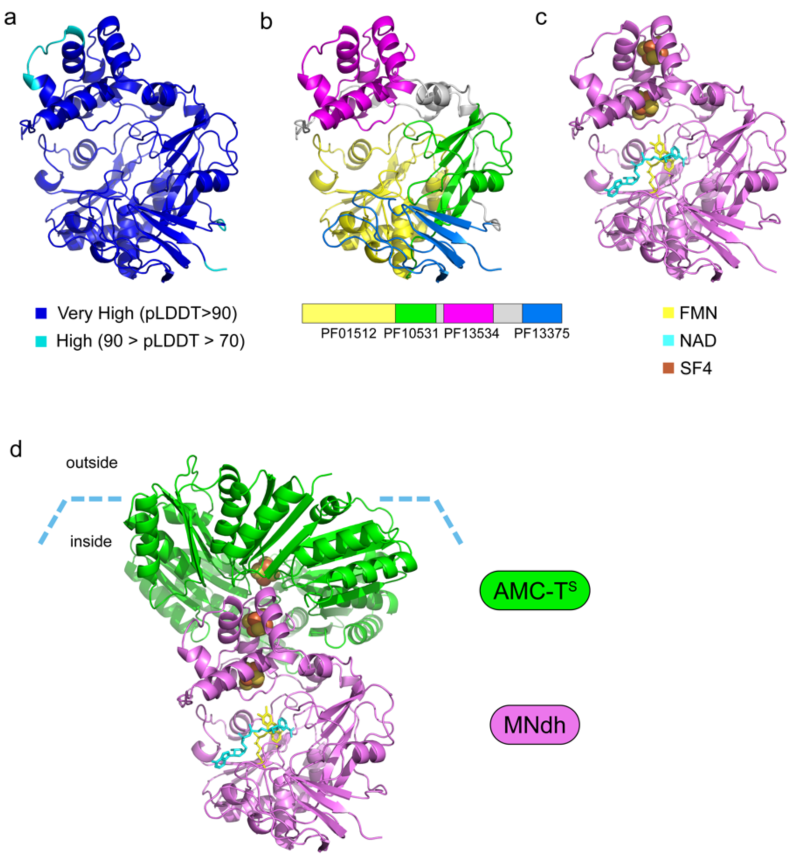


**Supplementary Fig. 16 | Structural models of the MNdh and the MNdh-AMC-TSE complex from a Hodarchaeales genome (accession number: GCA_003144275) by AlphaFold3.**
a. AlphaFold3 confidence. b. Domains of MNdh. c. Cofactor and metal placement in the MNdh structure. d. Model of a MNdh-AMC-T$^{SE}$ complex.

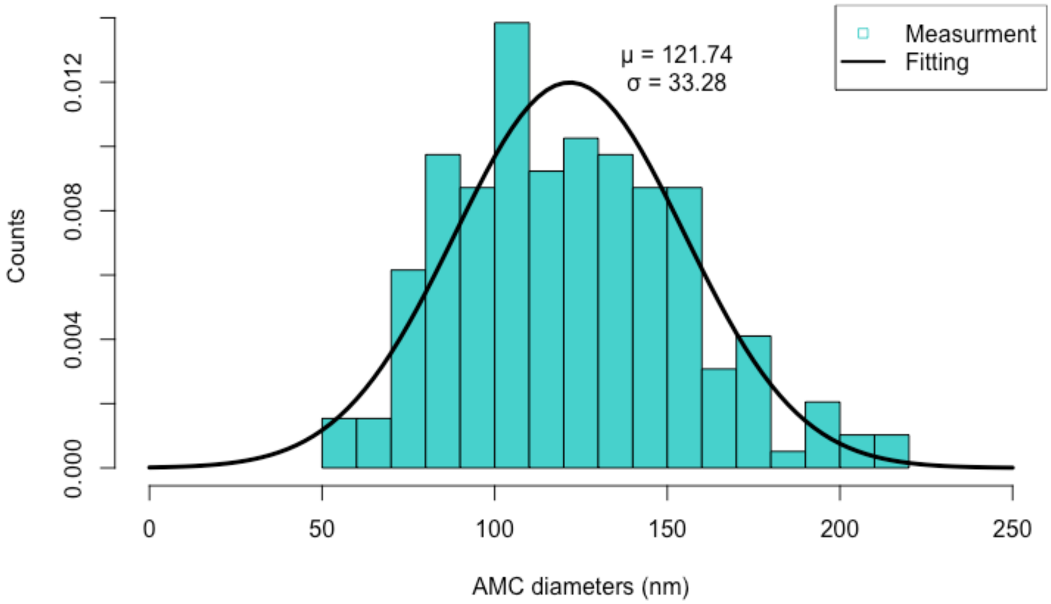


**Supplementary Fig. 17 | Distribution of AMC diameters: histogram and Gaussian probability density function fit.**

a

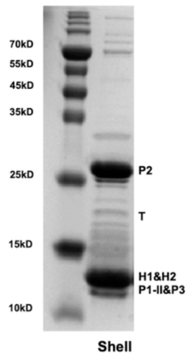


b

| Shell Protein | Score | Mass | Sequences | Coverage | emPAI |
|---|---|---|---|---|---|
| **P1-II** | 20332 | 11744 | 11 | 95.4% | 606.48 |
| **P3** | 7734 | 11621 | 11 | 88.8% | 294.9 |
| **H1** | 581 | 11136 | 6 | 81.6% | 10.42 |
| **P2** | 9437 | 20323 | 17 | 75.5% | 287.98 |
| **T** | 1090 | 18608 | 8 | 54.7% | 31.78 |

**Supplementary Fig. 18 | SDS-PAGE analysis of in vivo assembled AMC shells (a) and the mass spectrometry result (b).**

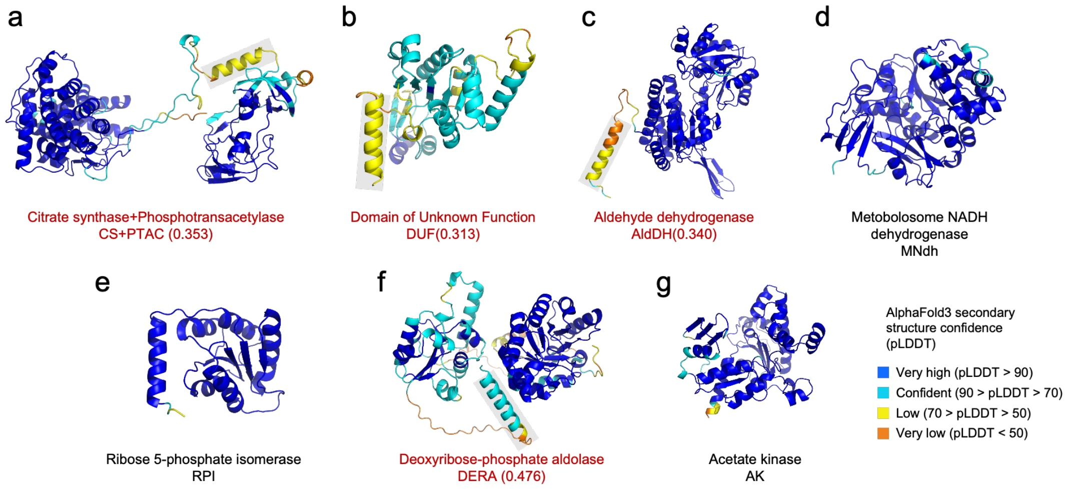


**Supplementary Fig. 19 | Structural models of cargo proteins from a Hodarchaeales genome (accession number: GCA_003144275).** a-g. AlphaFold3 model of metabolic enzymes colored by AlphaFold3 secondary structure confidence (pLDDT) (Blue = very high, cyan = confident, yellow = low, orange = very low). The encapsulation peptides (EPs) are highlighted with a gray box (the values in parentheses are the amphipathicity score[4]).

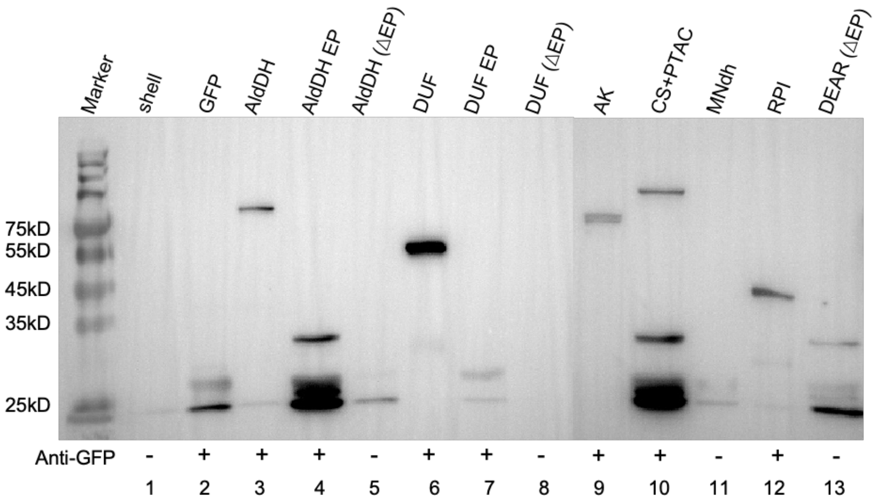


**Supplementary Fig. 20 | Western blot analysis of shell-cargo protein interactions mediated by encapsulation peptides (EPs).** To validate the function of EPs, Strep-II-tagged shell proteins were co-expressed with various GFP-tagged cargo proteins and isolated via Strep-tag affinity purification. Co-purified cargo was detected by Western blot using an aniti-GFP antibody. A GFP signal indicates a direct interaction between the shell particle and the cargo. Lane 1: Purified shell protein only (control). Lane 2: GFP only (control). Lanes 3–13: Co-expression of shell proteins with the following GFP tagged cargo proteins: full-length acetaldehyde dehydrogenase (AldDH)-GFP, the AldDH EP-GFP, truncated AldDH(ΔEP)-GFP, a full-length domain of unknown function (DUF)-GFP, the DUF EP-GFP, truncated DUF(ΔEP)-GFP, acetate kinase (AK)-GFP, citrate synthetase (CS)+phosphotransacetylase (PTAC)-GFP, metabolosome NADH dehydrogenase (MNdh)-GFP, ribose 5-phosphate isomerase (RPI)-GFP, and truncated deoxyribose 5-phosphate aldolase (DEAR(ΔEP))-GFP, respectively.

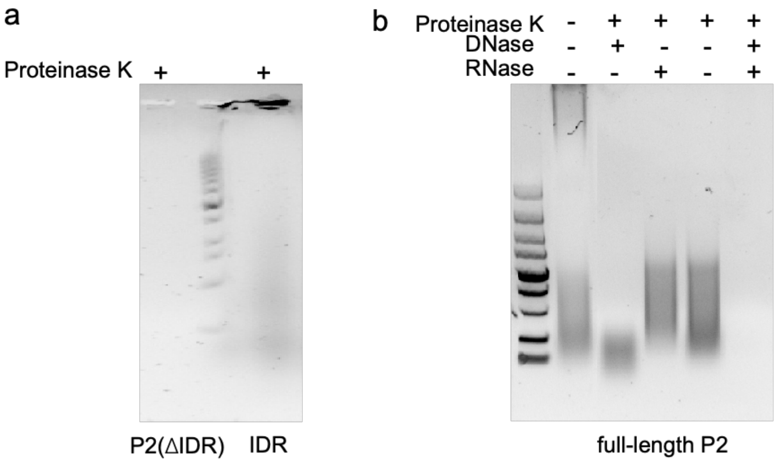


**Supplementary Fig. 21 | The AMC-P2 intrinsically disordered region (IDR)-nucleic acid binding test. a.** Agarose gel analysis of proteinase K-treated lysates from cells expressing either a truncated AMC-P2 protein (lacking the IDR) or the isolated IDR. No bands remained for the truncated AMC-P2, whereas obvious bands remained for the IDR. **b.** Migration of the full-length AMC-P2 expressed in vivo on an agarose gel after treatment with different enzymes. Lane 1: Untreated; Lane 2: Treated with Proteinase K and DNase; Lane 3: Treated with Proteinase K and RNase; Lane 4: Treated with Proteinase K; Lane 5: Treated with Proteinase K, DNase, and RNase

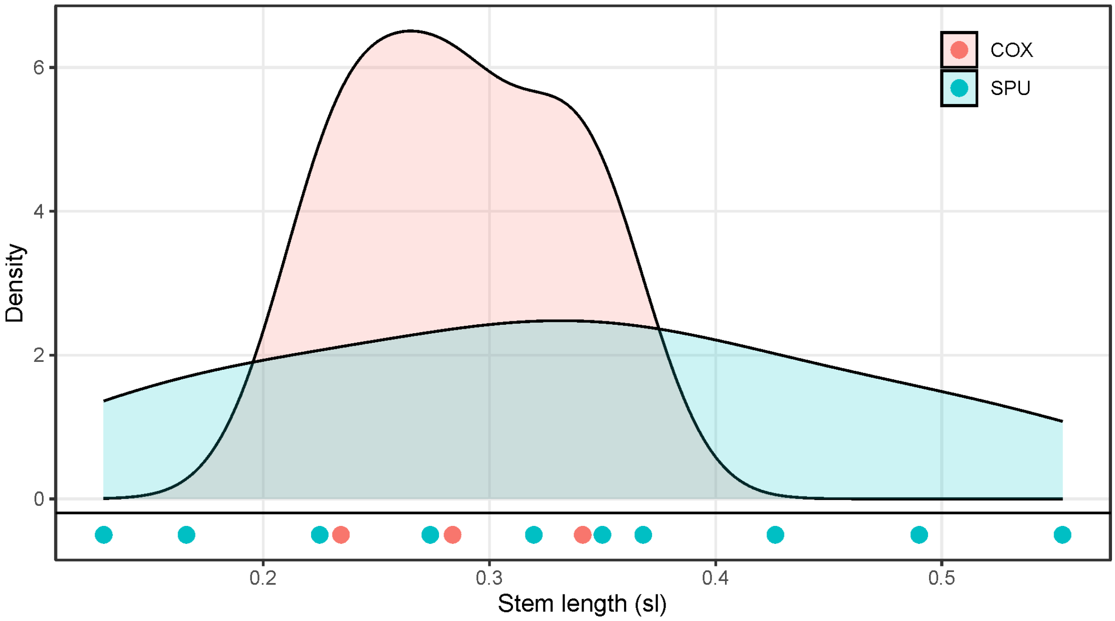


**Supplementary Fig. 22 | Stem length analysis for AMC and COX genes within Hodarchaeales.** Stem length (sl) was determined as the raw stem length of Hodarchaeales divided by the median of distance from terminal nodes to the last Hodarchaeales common ancestor (LHoCA). Stem length values of each AMC gene and COX subunit are shown using blue and red dots, respectively, and summarized into density plot.

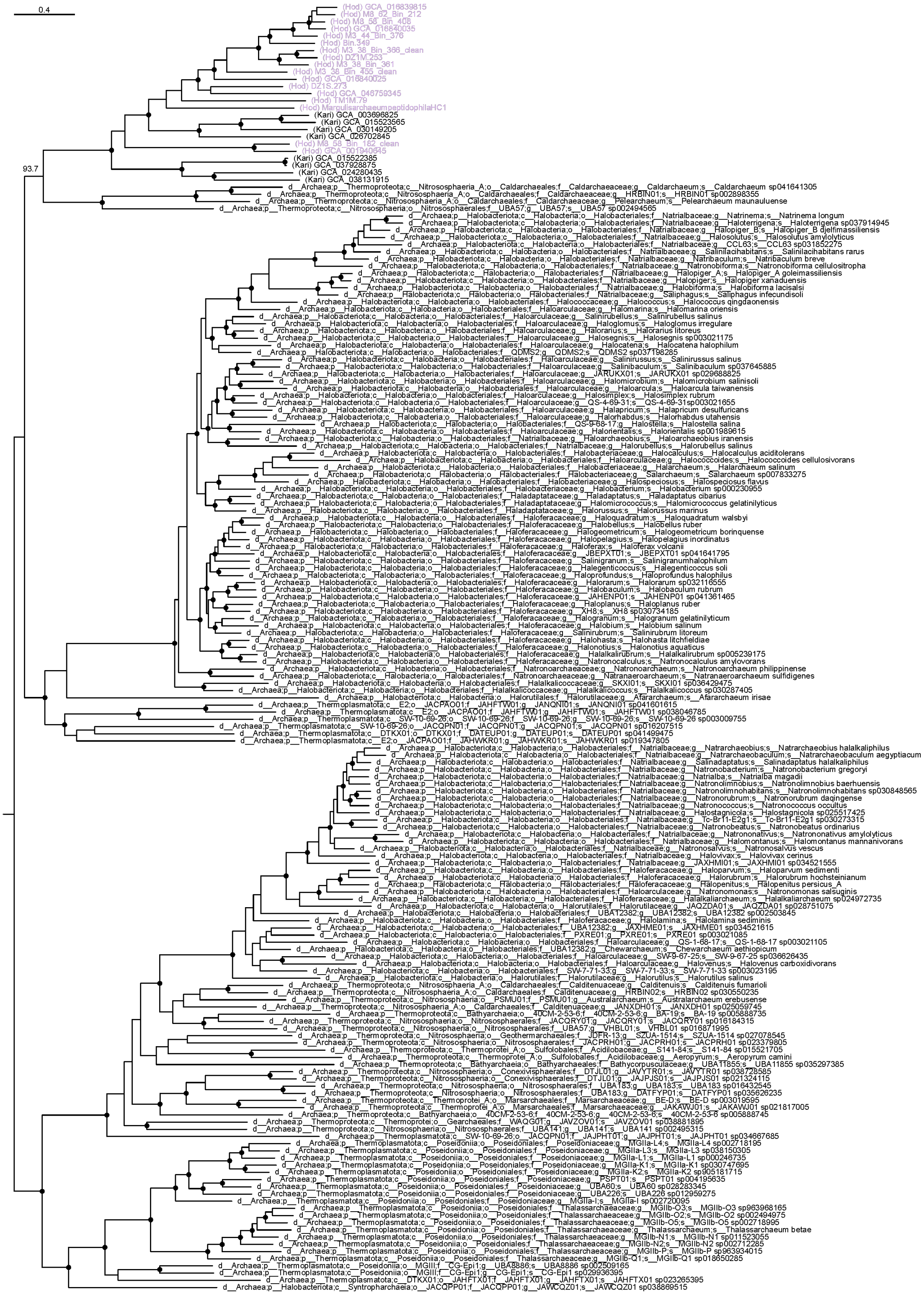

**Supplementary Fig. 23 | Phylogenetic analysis of COX genes.** The phylogenetic tree was constructed using IQ-tree with LG+C60+F+G model and posterior mean site frequency (PMSF) approximation, based on concatenated alignments of three subunits of the COX gene (coxABC, 1597 amino acids in total). Rooting position was selected using the minimal ancestor deviation (MAD) method. Nodes with SH-aLRT bootstrap values greater than 95% are indicated by solid circles. Hodarchaeales sequences are shown in light purple.
